# Structure and DNA bridging activity of the essential Rec114–Mei4 trimer interface

**DOI:** 10.1101/2023.01.18.524603

**Authors:** Kaixian Liu, Emily M. Grasso, Stephen Pu, Shixin Liu, David Eliezer, Scott Keeney

## Abstract

The DNA double-strand breaks (DSBs) that initiate meiotic recombination are formed by an evolutionarily conserved suite of factors that includes Rec114 and Mei4 (RM), which regulate DSB formation both spatially and temporally. *In vivo*, these proteins form large immunostaining foci that are integrated with higher order chromosome structures. *In vitro*, they form a 2:1 heterotrimeric complex that binds cooperatively to DNA to form large, dynamic condensates. However, understanding of the atomic structures and dynamic DNA binding properties of RM complexes is lacking. Here, we report a structural model of a heterotrimeric complex of the C-terminus of Rec114 with the N-terminus of Mei4, supported by nuclear magnetic resonance experiments. This minimal complex, which lacks the predicted intrinsically disordered region of Rec114, is sufficient to bind DNA and form condensates. Single-molecule experiments reveal that the minimal complex can bridge two or more DNA duplexes and can generate force to condense DNA through long-range interactions. AlphaFold2 predicts similar structural models for RM orthologs across diverse taxa despite their low degree of sequence similarity. These findings provide insight into the conserved networks of protein-protein and protein-DNA interactions that enable condensate formation and promote formation of meiotic DSBs.

## Introduction

Homologous recombination during meiosis promotes accurate chromosome segregation and genetic diversification in most sexually reproducing organisms. Meiotic recombination starts with DNA double-strand breaks (DSBs) formed by Spo11 protein (related to archaeal topoisomerase VI) plus a cohort of additional conserved factors (Keeney, 2008; Robert et al., 2016). Among these factors are Rec114 and Mei4, which are essential for DSB formation but also regulate the number, timing, and location of DSBs in many species (Henderson et al., 2006; Carballo et al., 2013; Rosu et al., 2013; Stamper et al., 2013; Murakami and Keeney, 2014; Kumar et al., 2010, 2018; Papanikos et al., 2019; Boekhout et al., 2019; Mu et al., 2020; Hinman et al., 2021; Claeys Bouuaert et al., 2021).

Rec114 and Mei4 from *Saccharomyces cerevisiae* form a 2:1 heterotrimeric complex *in vitro* and assemble cooperatively on DNA to form dynamic nucleoprotein condensates (Claeys Bouuaert et al., 2021a; Yadav and Claeys Bouuaert, 2021). *In vivo*, they associate early in meiotic prophase I with chromatin and form colocalized and interdependent foci along chromosome axes in multiple species (Li et al., 2006; Maleki et al., 2007; Panizza et al., 2011; Rosu et al., 2013; Stamper et al., 2013; Kumar et al., 2010, 2018; Boekhout et al., 2019; Papanikos et al., 2019; Hinman et al., 2021). Rec114–Mei4 (RM) complexes from different organisms interact directly with the meiotic TopoVI-like complex (Arora et al., 2004; Maleki et al., 2007; Claeys Bouuaert et al., 2021; Hinman et al., 2021; Vrielynck et al., 2021; Nore et al., 2022), but the molecular mechanisms of RM function remain poorly understood.

Rec114 and Mei4 were first recognized to function as a unit in *S. cerevisiae* (Arora et al., 2004; Li et al., 2006; Maleki et al., 2007). Their homologs in non-fungal species were not identified until later because of poor sequence conservation (Kumar et al., 2010). The Rec114 N terminus contains six signature sequence motifs (SSMs) defined by remote homology detection, plus a seventh SSM near the C terminus following a region of predicted disorder (**Fig. 1a**) (Maleki et al., 2007; Kumar et al., 2010; Tessé et al., 2017). X-ray crystallography of a fragment of mouse REC114 showed that the N-terminal SSMs correspond to diverse secondary structure elements within a pleckstrin homology (PH) domain (Kumar et al., 2018; Boekhout et al., 2019) that interacts with SPO11 partner TOP6BL and with vertebrate-specific DSB regulator ANKRD31 (Boekhout et al., 2019; Nore et al., 2022). The C-terminal region of yeast Rec114 (including SSM7) is sufficient to form a trimeric complex with the N terminus of Mei4 (including the first two of Mei4’s six SSMs), and this minimal complex is sufficient to bind DNA in an apparently sequence-nonspecific manner (Claeys Bouuaert et al., 2021a).

**Fig. 1:**
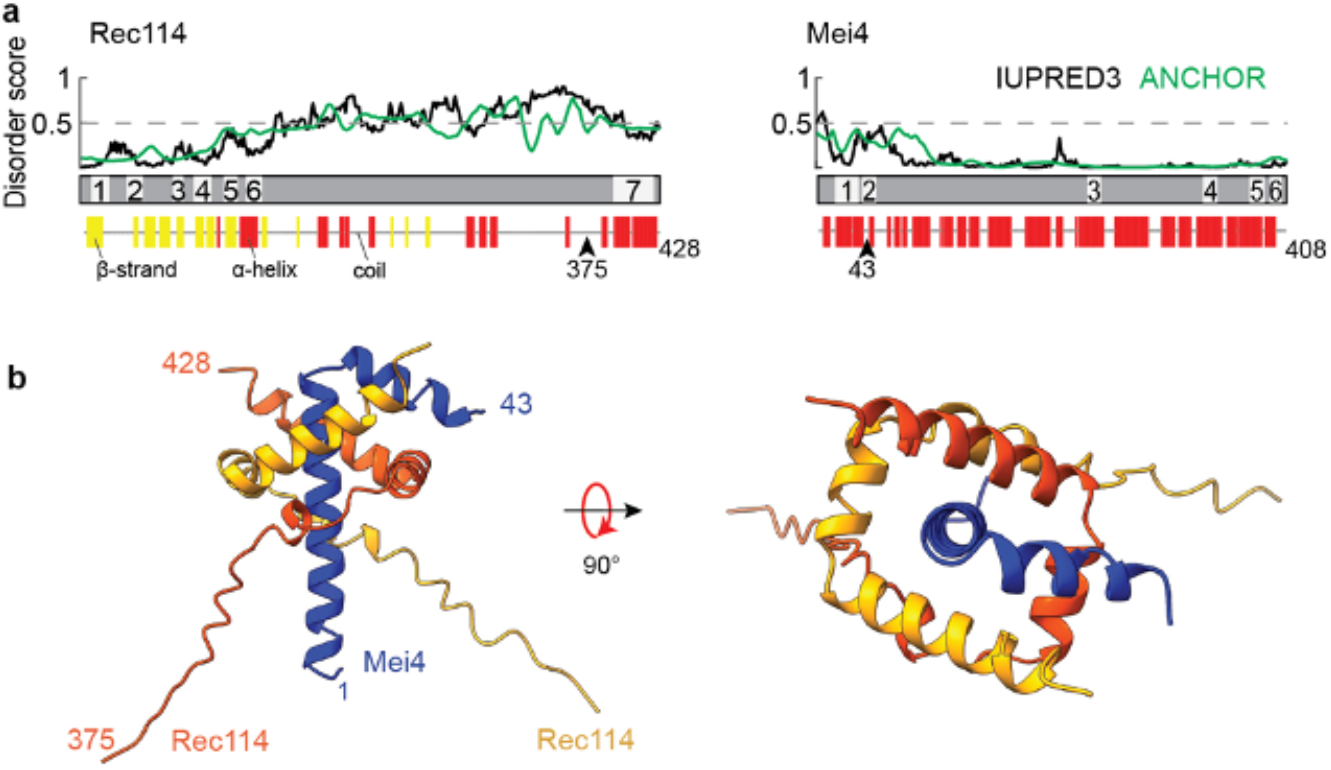
Structural prediction for trimers of Rec114_C_ and Mei4_N_. (a) Sequence-based secondary structure and disorder predictions (Methods) suggest that the C terminus of Rec114 and N terminus of Mei4 are ordered and primarily α-helical. Numbered segments are the SSMs from Kumar et al. (2010). (b) AlphaFold2 structure prediction for a heterotrimer of Rec114_C_ and Mei4_N_.

Aside from the structure of the mouse REC114 PH domain, there is little empirical information about the molecular structures of these essential, conserved meiotic DSB factors. Understanding is also limited about the DNA binding activities that support cooperative assembly of RM condensates. To address these issues, we examined the structures and biophysical properties of Rec114 and Mei4 using a combination of computational modeling, NMR spectroscopy, and bulk biochemical and singlemolecule experiments. We demonstrate an evolutionarily conserved structure for the RM trimerization and DNA binding (TDB) domain, and show that this minimal domain is sufficient to bind cooperatively to DNA to form nucleoprotein condensates. We further uncover a DNA-bridging activity of the RM-TDB domain that can bundle coaligned DNA molecules.

## Results

### Predicted structure of the Rec114–Mei4 trimer interface

On the basis of crosslinking plus mass spectrometry, yeast two-hybrid analyses, and purification of truncated recombinant proteins expressed in *Escherichia coli*, we previously showed that residues 375–428 of yeast Rec114 (hereafter Rec114_C_) and 1–43 of Mei4 (Mei4_N_) form stable trimers (Maleki et al., 2007; Claeys Bouuaert et al., 2021). Armed with this information, we used AlphaFold2 (Jumper et al., 2021; Mirdita et al., 2022) to predict a structure of this complex (**Fig. 1b and Fig. s1a,b**). Residues 389–426 from each Rec114 segment are predicted to form a twisted U shape consisting of three a helices (residues ~389–396, 399–407, and 409–426), with the two copies interlocking like a scissor staircase. Mei4 residues 3–42 are predicted to form an L-shaped helixturn-helix: α-helix 1 (residues 3–29) is embraced by the Rec114 dimer and lies along the dimer’s axis of rotational symmetry, while the shorter Mei4 α-helix 2 (31–42) lies across α-helix 2 of one of the Rec114 protomers. DALI searches (Holm and Laakso, 2016) revealed no matches to this structure.

The Rec114 dimer by itself is rotationally symmetric in the model (RMSD 0.6 Å for superimposition of the two copies of Rec114_389–426_; **Fig. s1c**). Mei4 breaks this symmetry because different faces of its first helix interact with the two Rec114 copies and because its second a helix contacts only one of the Rec114 protomers (**Fig. 1b**). AlphaFold2 did not generate a high-confidence prediction for Rec114 residues 375–388, possibly indicating that these are disordered (**Fig. s1b**).

### Experimental validation of the computational structure model

We tested the AlphaFold2 model empirically by examining recombinant Rec114_C_–Mei4_N_ complexes purified after co-expression in *E. coli* (**Fig. 2a**). Mei4_N_ could not be purified separately, precluding reconstitution of the complex from separately expressed components. The far-UV circular dichroism (CD) spectrum of Rec114_C_–Mei4_N_ trimers showed characteristic minima at 208 and 222 nm typical for α-helical proteins (**Fig. 2b**) (Greenfield, 2007).

**Fig. 2:**
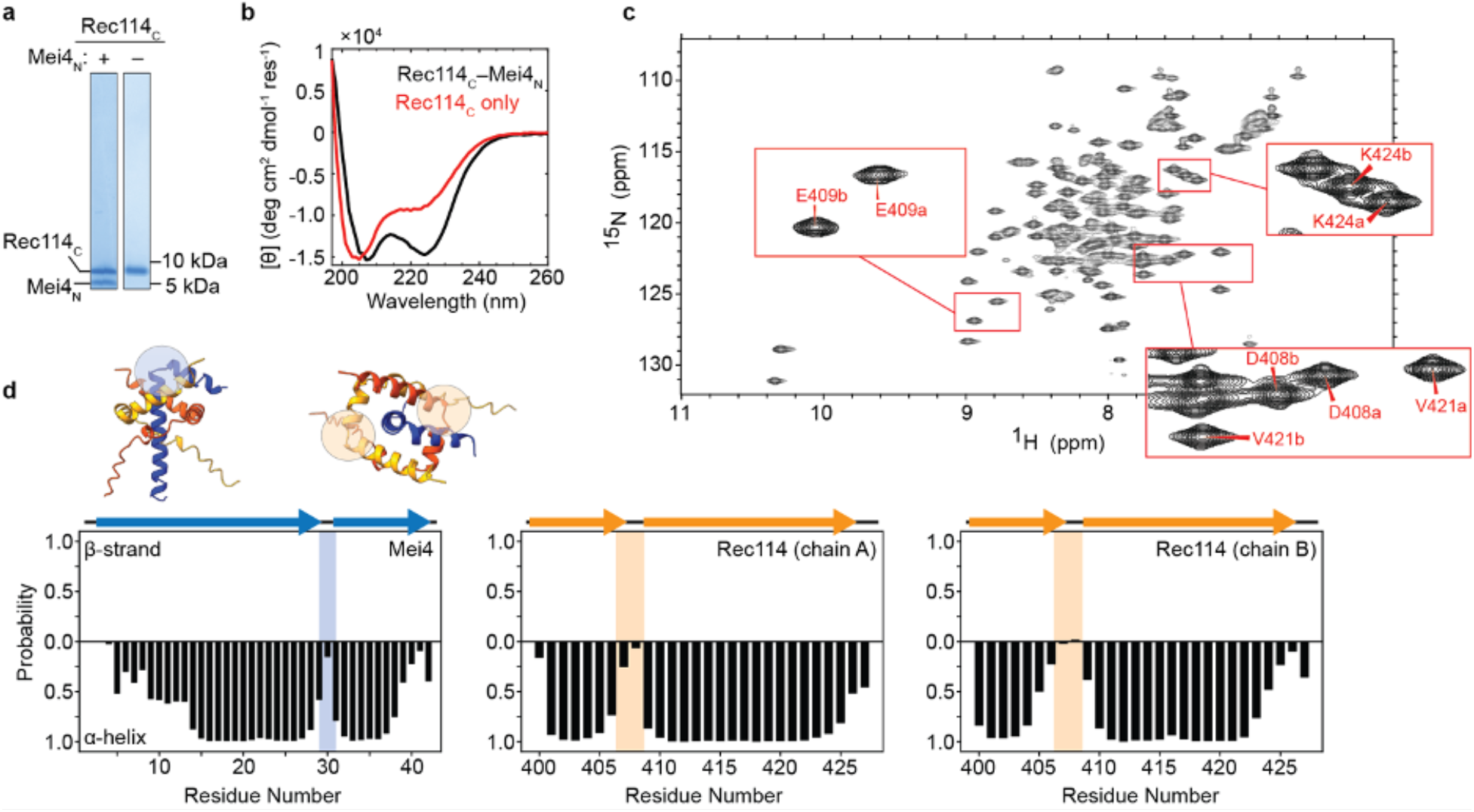
Spectroscopic analysis of Rec114_C_ complexes with Mei4_N_. (a) SDS-PAGE of purified proteins. (b) CD spectra for Rec114_C_ with or without coexpressed Mei4_N_. (c) Two-dimensional {^1^H-^15^N} HSQC spectrum of Rec114_C_–Mei4_N_ complexes. Insets show examples of distinct peaks assigned to cognate residues from the two Rec114_C_ chains. (d) TALOS-N secondary structure analysis confirming helix-turn-helix segments in Rec114_C_ and Mei4_N_. The arrows above the plots show positions of helices predicted by AlphaFold2. Shaded regions are the turns highlighted in the AlphaFold2 model (insets).

Solution nuclear magnetic resonance (NMR) spectroscopy experiments using the uniformly {^15^N-^13^C}-labeled ternary complex showed moderate peak dispersion in two-dimensional {^1^H-^15^N} heteronuclear single quantum coherence (HSQC) spectra, consistent with expectations for well-structured helical proteins (**Fig. 2c**). Backbone chemical shifts were assigned for residues 399–428 of Rec114_C_ and residues 5–42 of Mei4_N_ at pH 7.4 (**Fig. s2** and Methods). Importantly, Rec114_C_ exhibited two sets of peaks corresponding to the two copies in the trimeric complex (**Fig. 2c insets** and **Fig. s2a,b**), consistent with the predicted asymmetry between the Rec114_C_ protomers.

Secondary structure prediction from chemical shifts using TALOS-N (Shen and Bax, 2013) supported the AlphaFold2 predictions for helixturn-helix segments at residues 399–428 of both copies of Rec114_C_ (α-helices 2 and 3) and at residues 5–42 of Mei4_N_ (**Fig. 2d**). Notably, Mei4_N_ residues 5–13 were predicted to be fractionally helical, suggesting fraying at the N-terminal end of the first Mei4_N_ helix and consistent with the lower confidence of the AlphaFold2 prediction for this region (**Fig. s1a,b**). No NMR signals were observed for Rec114_C_ residues 386–398, precluding assessment of their structure. The TALOS-N predictions were corroborated by NOEs between consecutive amide protons in both Rec114_C_ and Mei4_N_. consistent with expectations for helical structure (**Fig. s2**).

Given the absence of NMR signals for Rec114_C_ residues 385–398 and the fractional helicity of Mei4_N_ residues 5–13, we examined the effects of truncating each construct. Removing 13 amino acids from the N-terminus of Rec114_C_ (Rec114_388-428_) resulted in minimal spectroscopic changes other than the elimination of a few peaks originating from the very N terminus of Rec114_C_ (**Fig. s3a**). In contrast, removing 24 residues (Rec114_399-428_) resulted in substantial chemical shift perturbations, the loss of several well-dispersed resonances, and broadened linewidths (**Fig. s3a**), consistent with the loss of well-defined structure in the complex. These truncations suggest that Rec114_C_ residues 388–398, but not residues 375–387, are important for structural stability. Removing the first 12 residues of Mei4_N_ (Mei4_13-43_) also resulted in minimal perturbations in spectra (**Fig. s3b**). Together, these data indicate that residues 388–428 of Rec114 and 13-43 of Mei4 form the core structured unit of the Rec114–Mei4 interface.

Given the importance of Rec114_C_ residues 388–398 to complex stability, we sought conditions under which we could observe NMR signals for this region. Lowering the pH, which slows amide proton exchange with solvent (Matthew and Richards, 1983), resulted in the appearance of a number of new peaks with minimal perturbations elsewhere in the spectra (**Fig. s3c**). Assignment of the minimal structured construct (Rec114_388-428_–Mei4_13-43_) at pH 6.1 revealed that these new peaks correspond to residues 388–399 of Rec114_C_ (**Fig. s4a,c,e**). TALOS-N secondary structure predictions for these constructs (**Fig. s4b,d,f**) showed that truncation of both Rec114_C_ and Mei4_N_ had little effect on helical structures that had been evident in the longer construct. Surprisingly, no stable secondary structure was predicted for the newly visible regions comprising residues 388–399. Although the AlphaFold2 model indicates α-helices for Rec114 protomers at residues 391–394 or 388–396, the confidence score for this prediction is low (**Fig. s1a,b**). Our data suggest that despite its importance for the stability of the complex, this region does not adopt stable helical structure in the absence of DNA at lower pH. Altogether, these spectroscopic data are in good agreement with the AlphaFold2 model.

Although the Rec114_C_ C-terminal region by itself can dimerize (Claeys Bouuaert et al., 2021a), the CD spectrum of purified Rec114_C_ alone showed substantially diminished α-helical character, indicating that it is less structured (**Fig. 2b**). In contrast to the ternary complex, HSQC spectra of Rec114_C_ alone showed poor dispersion, with a limited number of peaks of variable intensity (**Fig. s5a**). Purified Rec114_C_ also eluted as a broader peak compared with the ternary complex in size exclusion chromatography (**Fig. s5b**). These findings suggest that Rec114_C_ by itself is at least partly unfolded, suggesting in turn that interaction with Mei4_N_ stabilizes the α-helical secondary structure of Rec114.

### Structural insights into sequence conservation

The model accounts well for patterns of sequence conservation, with the structural motifs elucidated in the model and our experimental data corresponding to conserved elements. In Rec114, SSM7 comprises helices 2 and 3 plus the turn between them (**Fig. 3a**). In Mei4, SSM1 corresponds to the stable second half of helix 1 plus a part of helix 2, while SSM2 begins in helix 2 and extends two residues beyond the structured domain analyzed here (**Fig. 3a**).

**Fig. 3:**
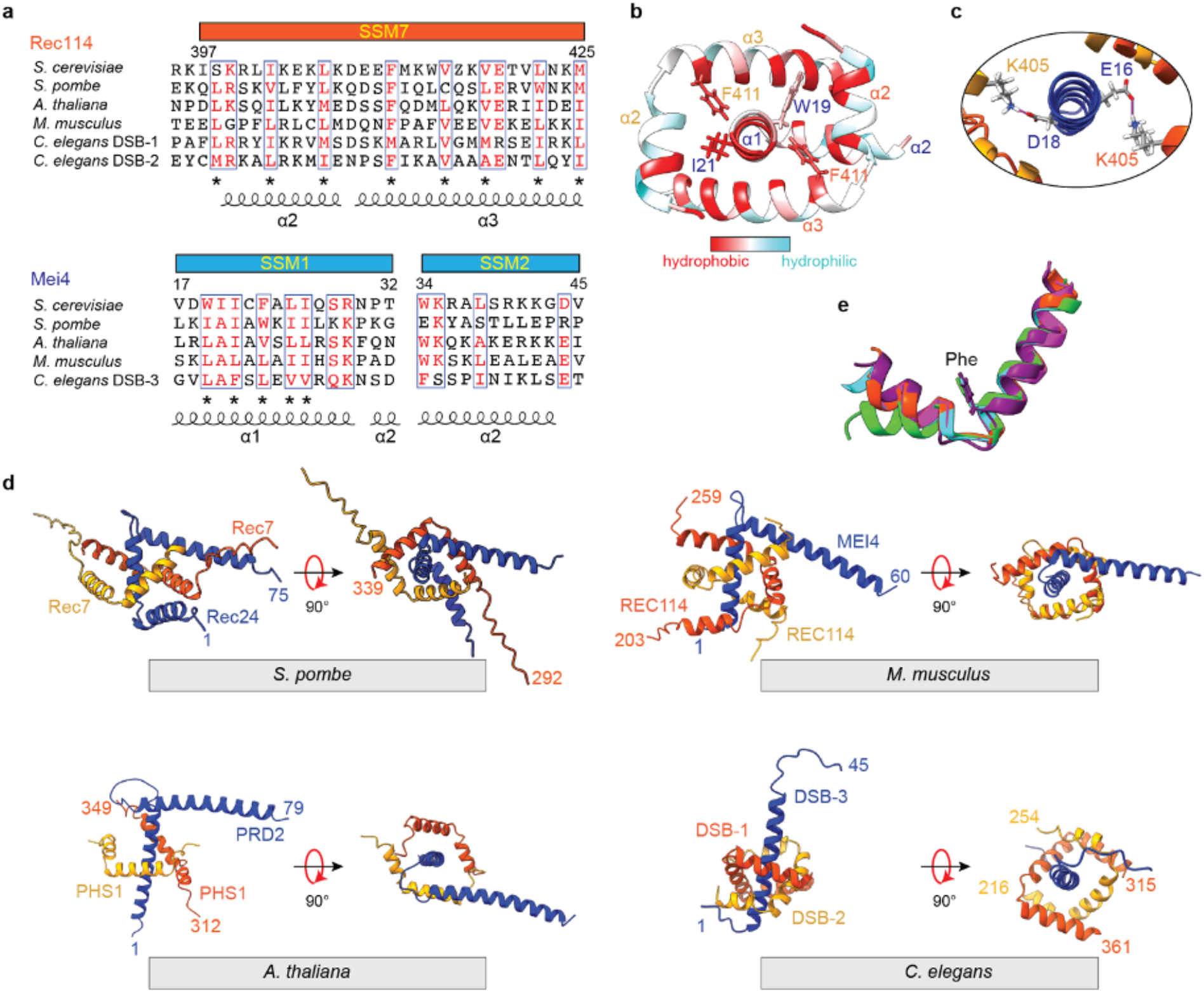
Structure and conservation of Rec114–Mei4 interactions. (a) Structure-informed alignment of Rec114 and Mei4 orthologs. Secondary structure elements and SSMs are indicated. Conserved residues are highlighted in red, and conserved hydrophobic residues are labeled by asterisks. (b) Hydrophobic interactions at the Rec114_C_–Mei4_N_ interface. Hydrophobic residues are highlighted in red. (c) Predicted contacts between K405 from each Rec114_C_ chain and either E16 or D18 from Mei4_N_. (d) AlphaFold2 models of equivalent domains for Rec114 and Mei4 orthologs from *S. pombe, M. musculus, A. thaliana, C. elegans*. (e) Structural alignment of SSM7 from Rec114 orthologs. The highly conserved phenylalanine is shown (F411 in *S. cerevisiae*). Orange red, *S. cerevisiae;* cyan, *S. pombe* Rec7; purple, *M. musculus* REC114; green, *A. thaliana* PHS1/AtREC114; magenta, *C. elegans* DSB-2.

Many highly conserved residues are hydrophobic and contribute to intermolecular interfaces in the model. F411 is nearly invariant in Rec114 orthologs (**Fig. 3a**) and contacts Mei4 residues W19 or I21 (**Fig. 3b**), the equivalents of which are universally large hydrophobic residues (**Fig. 3a**). Similarly, W34 of Mei4 helix 2 contacts I412 and V415 from one of the Rec114 chains (**Fig. s6a**). W34 is the first residue in SSM2 and is nearly invariant in Mei4 orthologs; V415 in Rec114 is also highly conserved, while I412 is more moderately conserved (**Fig. 3a**). Rec114 helices 2 and 3 are amphipathic, with conserved hydrophobic side chains every 3–4 residues facing inward towards Mei4 (**Fig. 3a,b**). In contrast, the conserved hydrophobic residues in helix 1 of Mei4 are not restricted to just one face, appearing every 1–2 residues (**Fig. 3a,b**), consistent with this helix being embraced by the two copies of Rec114. The model also predicts a salt bridge between the nearly invariant E419 of Rec114 and R29 in Mei4, which is nearly always a basic residue (**Fig. 3a and Fig. s6a**).

Two-fold symmetric hydrophobic contacts between the Rec114 chains also contribute to the sequence conservation. I402 and L406 from helix 2 of each Rec114 copy pack against hydrophobic residues V418, L422, and M425 from helix 3 of the other chain (**Fig. s6b**). These residues are highly conserved (**Fig. 3a**).

There are also a number of predicted interactions involving residues that are less well-conserved, including salt bridges between Mei4 E16 and D18 and K405 from each Rec114 chain (**Fig. 3c**) and hydrogen bonds between Mei4 K41 and backbone carbonyl oxygens from L406 and K407 of one copy of Rec114 (**Fig. s5a**).

To further explore the correspondence between sequence conservation and structure, we generated AlphaFold2 models for trimeric RM complexes from *Schizosaccharomyces pombe, Mus musculus, Arabidopsis thaliana*, and *Caenorhabditis elegans* (**Fig. 3d and Fig. s6c**). For *C. elegans*, we used the two Rec114 paralogs, DSB-1 and DSB-2 (Rosu et al., 2013; Stamper et al., 2013). Consistent with an independent analysis (Guo et al., 2022), the overall folds were similar and predicted with high confidence scores: two copies of Rec114 form an approximately two-fold symmetric interlocking set of helix-turn-helix motifs embracing an a helix from Mei4. Alignment of the models for Rec114 SSM7 illustrates the strong conservation of the helix boundaries and the relative orientation between helices 2 and 3 (**Fig. 3e**). The nearly invariant phenylalanine (budding yeast F411) occupies the same position after the turn between these helices, highlighting its conserved role in Rec114–Mei4 complex formation. Additionally, the R29–E419 salt bridge is conserved in most of the predicted structures other than plants (although this may be due to the low confidence of that region in the predicted structure of plants (**Fig. s6c**)), and this salt bridge in worms is formed between DSB-2^Rec114^ and DSB-3^Mei4^, but would not be able to form with DSB-1^Rec114^, which has an arginine at the equivalent position to E419 (**Fig. 3a**).

Although the overall folds were similar, there were also substantial differences, in keeping with the high degree of sequence variability between species that has been previously described (Keeney, 2008; Kumar et al., 2010; Tessé et al., 2017). For example, the trajectory of the Mei4 turn and second helix and the nature of the interaction of that second helix with Rec114 is markedly different between species (**Fig. 3d**). Additionally, AlphaFold2 predicted an extra helix after SSM7 in both DSB-1 and −2 from *C. elegans*, but not in other Rec114 orthologs examined (**Fig. 3d**). Therefore, unlike mouse and budding yeast Rec114 that use an N-terminal helix, the *C. elegans* Rec114 orthologs use a C-terminal helix to form a U-shaped helical pocket (**Fig. 3d**). Also, the poor prediction confidence of the DSB-3 SSM2 region (pLDDT score in **Fig. s6c**) is consistent with this segment being less well conserved (Hinman et al., 2021).

### The RM-TDB domain is sufficient to form condensates with DNA

Rec114_C_–Mei4_N_ trimers are competent to bind pUC19 plasmid DNA substrates, but under the electrophoretic mobility shift assay (EMSA) conditions tested, they did not appear to be able to form condensates (Claeys Bouuaert et al., 2021a). We therefore more fully characterized the DNA-binding activity of Rec114_C_–Mei4_N_ trimers (hereafter the RM-TDB domain, for “trimerization and DNA-binding”).

In EMSAs with a 150-bp substrate, we observed at least two discrete shifted species plus material trapped in the wells (**Fig. 4a**), indicating that multiple protein complexes could bind to the same or multiple copies of DNA. We cannot accurately estimate K_d_ values because we do not know the stoichiometry of protein bound to DNA or the number of binding sites per DNA molecule, so we compared binding to different substrates by measuring the protein concentration that resulted in 50% of the DNA being bound (C_50_). The RM-TDB domain showed roughly comparable abilities to bind to linear DNA substrates of different lengths ranging from 80 to 1000 bp (C_50_ of ~80–100 nM), while binding to a 20-bp substrate occurred with substantially lower apparent affinity (C_50_ of ~800 nM**; Fig. 4a,b and Fig. s7a,b**).

**Fig. 4:**
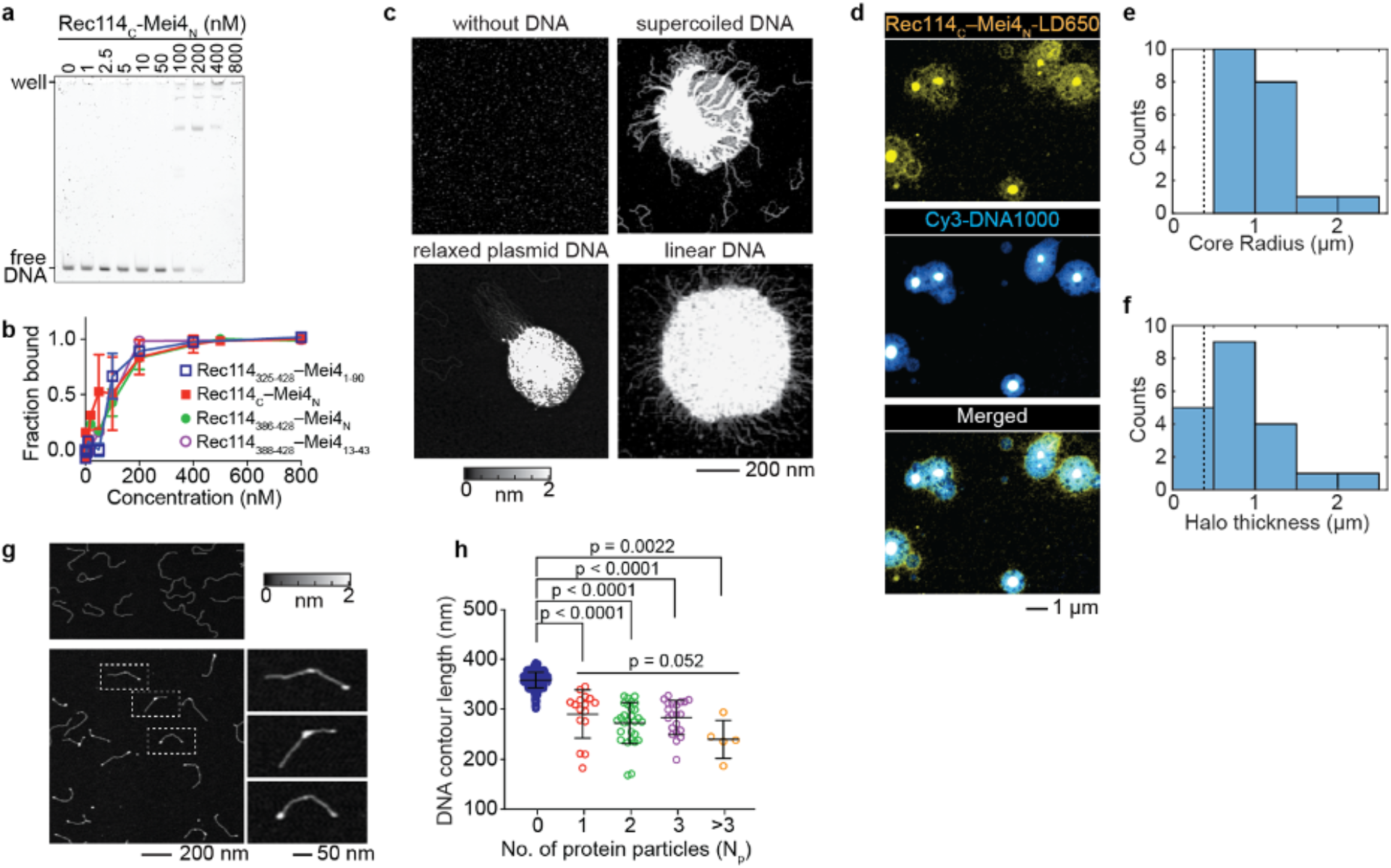
DNA binding and nucleoprotein condensate formation by the RM-TDB domain. (a) Representative EMSA of binding to a 150-bp DNA substrate. (b) Comparison of DNA binding (150-bp substrate) by RM-TDB complexes composed of different fragments of Rec114 and Mei4. Error bars indicate mean ± range (two replicate experiments) or mean ± SD (three replicates). C_50_ values were 90 ± 20 nM (Rec114_N_–Mei4_1-90_, mean ± range); 90 ± 30 nM (Rec114_C_–Mei4_N_, mean ± SD); 110 ± 30 nM (Rec114_386-428_–Mei4_N_, mean ± range). The EMSA for Rec114_388-428_–Mei4_13-43_ was conducted once (C_50_ of ~100 nM). (c) AFM images of 200 nM Rec114_C_–Mei4_N_ in the absence of DNA or presence of 1 ng/μl supercoiled pUC19 plasmid DNA, relaxed circular pHOT1 plasmid DNA, or 1000-bp linear DNA. (d) Confocal images of condensates formed by 450 nM fluorescently labeled (LD650) Rec114_C_–Mei4_N_ and 25 nM 1000-bp Cy3-DNA. (e, f) Quantification of the radius of central cores (panel e) and thickness of haloes (panel f) of the condensates from confocal images (N = 20). Dashed lines indicate the expected contour length of free DNA (0.383 μm). (g) AFM images of 1 ng/μl 1000-bp linear DNA in the absence (top) or presence (bottom) of 70 nM Rec114_C_–Mei4_N_. Examples of DNA molecules with bound protein (dashed boxes) are shown in zoomed insets to the right. (h) DNA contour lengths of free DNA (blue points) and protein-bound DNA (with N_p_ indicating the number of protein particles per DNA molecule). Error bars indicate mean ± SD (free DNA, 358 ± 16 nm (N = 474 DNA molecules); N_p_ = 1 particle bound, 290 ± 48 nm (N = 16 DNA molecules); N_p_ = 2, 272 ± 41 nm (N = 28 DNA molecules); N_p_ = 3, 283 ± 34 nm (N = 21 DNA molecules); N_p_ > 3, 240 ± 38 nm (N = 5 DNA molecules). The pairwise p values are from unpaired two-tailed Student’s t-tests. The group p value for different numbers of protein particles bound is from a Kruskal-Wallis test.

The apparent affinity for the 150-bp substrate was affected only modestly if at all by including 50 additional amino acids from the IDR of Rec114 and 47 additional residues from Mei4 (complexes of Rec114_325–428_ with Mei4_1–90_) (**Fig. 4b and Fig. s7c,d**). Binding was similar or identical with constructs lacking the structurally dispensable N-terminal residues from the Rec114 fragment and Mei4 (complexes of Rec114_388–428_ with Mei4_13–43_) (**Fig. 4b**). We conclude that the minimal folded RM-TDB alone is sufficient to bind DNA, albeit with substantially lower affinity than the full-length RM complex, which has C_50_ of 6 nM for 80-bp DNA (Claeys Bouuaert et al., 2021a).

We visualized protein-DNA complexes using atomic force microscopy (AFM) and fluorescence confocal microscopy. At a concentration of the RM-TDB domain (200 nM) above the C_50_ and in the presence of supercoiled or relaxed circular plasmid DNA or 1000-bp linear DNA, AFM showed large clusters with DNA emanating out from a dense core (**Fig. 4c**). In contrast, the protein alone at concentrations ranging from 200 nM to 2 μM formed only small, relatively homogeneous particles on the mica surface (**Fig. 4c and Fig. s7e**). Clusters still formed in constructs without residues 375–387 from Rec114_C_ (**Fig. s7f**).

Interestingly, although Rec114_C_ alone bound to DNA with lower apparent affinity than the RM-TDB domain (C_50_ in EMSAs of ~300–400 nM; **Fig. s7g**), it could still form nucleoprotein clusters in AFM experiments at high concentration (**Fig. s7h**). Mei4_N_ may contribute directly to DNA binding and cooperative assembly, or may act primarily by stabilizing the Rec114_C_ fold.

When nucleoprotein assemblies were imaged by confocal microscopy of fluorescent RM-TDB domain and 1000-bp linear DNA, we observed dense protein- and DNA-rich cores surrounded by halos that also contained both protein and DNA, but at lower density (**Fig. 4d**). The dense cores (minimum radius 580 nm; median 940 nm) were larger than the expected contour length of the DNA (383 nm) (**Fig. 4e**), indicating that many copies of the DNA are interconnected to form the cores. The thickness of the halos (median 657 nm) also typically exceeded the DNA contour length (**Fig. 4f**), ruling out a simple model that the halos consist solely of DNA molecules that have one end embedded in the core. Instead, we infer that the halos are also complex networks of protein-bound DNA molecules, some of which are embedded in the core and some of which are not. We note that these structures were imaged after immobilization on a surface but were formed in solution, so these are likely to be two-dimensional deformations of three-dimensional—presumably globular—structures.

These findings show that the minimal folded RM-TDB domain by itself is capable of assembling cooperatively with DNA to form large structures reminiscent of the nucleoprotein condensates formed by fulllength RM complexes (Claeys Bouuaert et al., 2021a). The critical concentration needed is considerably lower for the full-length proteins, however, which form condensates at RM concentrations as low as 12 nM under otherwise similar conditions. Thus, while the regions of the proteins outside of the RM-TDB are not strictly required, they clearly contribute to the efficiency of condensation.

We also examined DNA binding by the RM-TDB domain at a lower concentration (70 nM). No condensates were observed by AFM, but instead we found numerous discrete particles that were located both at DNA ends and interstitially (**Fig. 4g and Fig. s7i**). Both types of binding event yielded similarly sized particles as well (**Fig. s7j**). The contour length of the 1000-bp substrate was markedly shorter when bound by protein (**Fig. 4h**), suggesting that binding of the RM-TDB domain compacts the DNA. By comparing the volumes of free and protein-bound DNA, we estimated that there were on average ~seven Rec114_C_–Mei4_N_ trimers per binding site (**Fig. s7k** and Methods).

### The RM-TDB domain reversibly bridges coaligned DNA molecules

To characterize the dynamics of DNA binding by the RM-TDB domain, we conducted single-molecule imaging experiments combining optical trapping with scanning confocal microscopy in a laminar-flow microfluidic system (Renger et al., 2021; Leicher et al., 2022) (**Fig. 5a**). RM-TDB was fluorescently labeled using the ybbR-Sfp system (Yin et al., 2006; Wasserman et al., 2019), in which the bacterial phosphopantetheinyl transferase Sfp covalently attached a single LD650 fluorophore to a specific serine residue in the 11-residue ybbR peptide fused to the C-terminus of Mei4_N_ (**Fig. s8a**). EMSA and NMR analyses indicated that the ybbR tag without the fluorophore does not affect the DNA-binding activity or structure of RM-TDB (**Fig. s8b,c**), and DNA binding was also unaffected by the labeling reaction (**Fig. s8b**). We used either a dual-trap or quadruple-trap system to capture one or two pairs of streptavidin-coated polystyrene beads to which biotinylated bacteriophage lambda DNA molecules could be bound and then moved into the channel containing a mixture of unlabeled and fluorescently labeled RM-TDB for visualization (**Fig. 5a**).

**Fig. 5:**
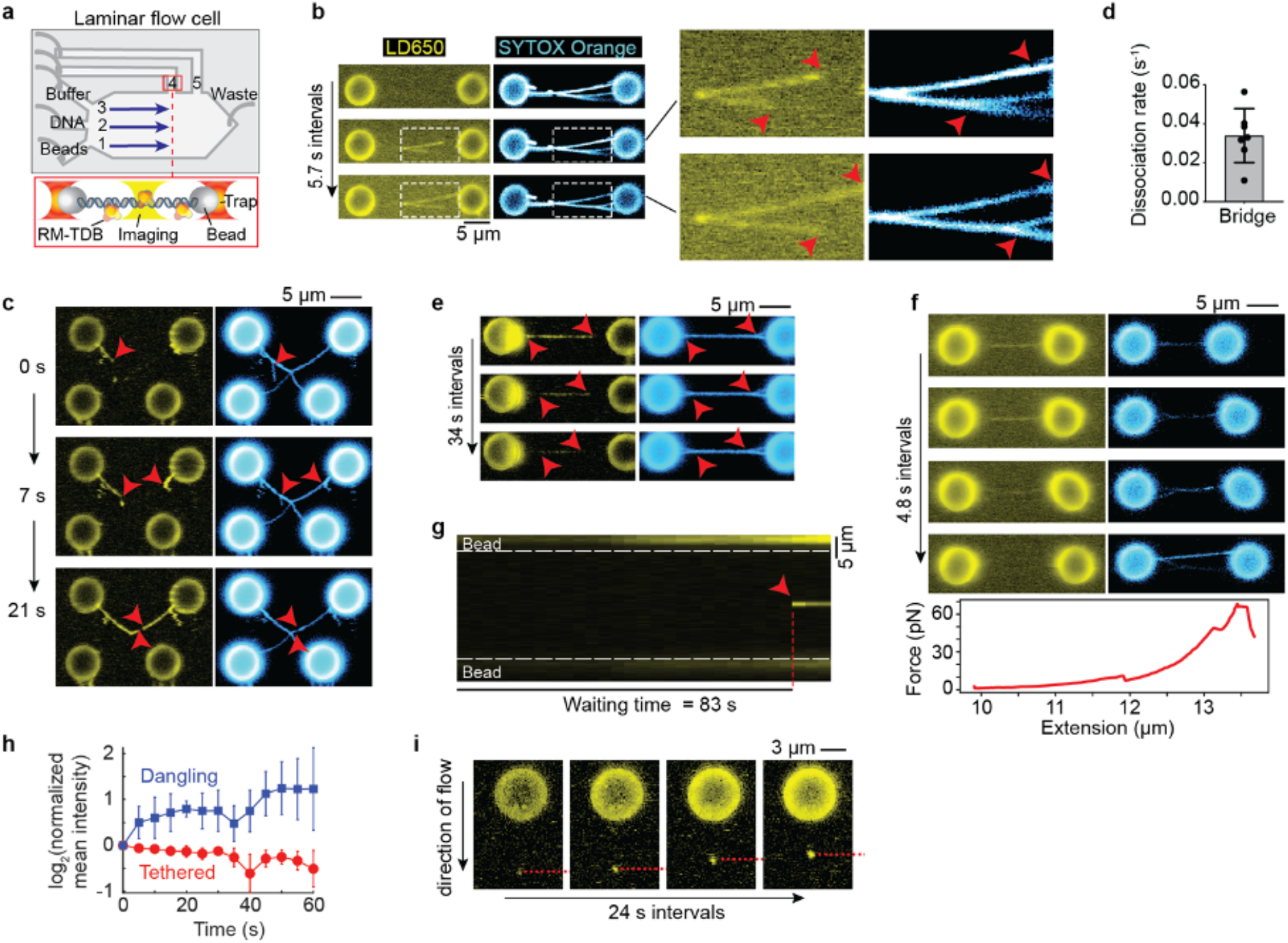
Single-molecule fluorescence imaging of interactions of the RM-TDB domain with DNA. (a) Schematic of experimental setup (not to scale). Streptavidin-coated beads, biotinylated *λ*-DNA and PBS buffer were separated by laminar flow in channel 1–3, respectively. After tether formation, beads were moved to channels 4 or 5 for protein loading and imaging. The RM-TDB protein concentration in all C-trap experiments was 20 nM. (b) Bridging of multiple tethered DNA duplexes by the RM-TDB domain. Arrows in insets indicate where separate DNA molecules branch apart, coinciding with the ends of RM-TDB tracks. See also Supplemental Movie 1. (c) Dangling DNA bundled together with stretched tethers. Arrows indicate dangling *λ*-DNA molecules (i.e., attached to only one bead) that are initially stretched out by flow but become progressively coaligned with segments from the tethers connecting the top pair of beads. Bundling of the dangling DNA with the tethers is coincident with extension of tracks of RM-TDB binding. Note that the single DNA tether that connects the lower pair of beads did not acquire any coating by the RM-TDB. See also Supplemental Movie 2. (d,e) Quantification of protein dissociation rates (panel d) and representative example (panel e) of disassembly of protein-DNA bridges moved into a protein-free channel. Red arrows in panel e indicate locations where the DNA molecules became separated. See also Supplemental Movie 3. Each point in panel d is a measurement from a single bridge (N=7; example in Fig. s9b); error bars indicate SD. (f) Force-promoted reversal of the RM-TDB bridge assembly. As beads connected by bridged tethers were pulled apart with increasing force, LD650 fluorescent signal decreased over time, indicating that RM-TDB was undergoing net dissociation despite free protein remaining available in the channel. Segments of the coaligned DNA tethers became separated coincident with loss of protein binding. The corresponding force-extension curve is plotted below. (g) Example kymograph of sudden focal binding of the RM-TDB domain (red arrow) to a stretched tether. The waiting time is the interval between introduction of the beads to the protein channel and first appearance of the focus. The white dashed lines indicate bead boundaries. See also Supplemental Movie 4. (h) Average change in protein fluorescence intensity over time for focal binding events on stretched tethers (red, N = 4) or on dangling DNA (blue, N = 9). The fluorescence signal at each time point was normalized to the signal in the first frame where binding of RM-TDB was detected (see Methods). Error bars indicate SD. (i) Accumulation of RM-TDB pulls dangling DNA against flow. A representative example is shown of RM-TDB binding to the tip of a dangling *λ*-DNA molecule bound to a single bead and stretched by flow. Over time, the protein-bound tip of the dangling DNA retracted upward toward the bead as indicated by the dashed red dashed line. See also Supplemental Movie 5.

We observed a striking DNA-binding activity for the RM-TDB domain when a pair of beads tethered together by two or more DNA molecules was moved into the protein channel. DNA segments that were already aligned were rapidly bound by RM-TDB complexes continuously along the length of the aligned regions (**Fig. 5b and Supplemental Movie 1**). With further incubation, additional RM-TDB protein associated with the DNA, simultaneously lengthening both the stretches of bound protein and the segments of aligned DNA (**Fig. 5b**).

DNA was not coated by protein when only a single lambda DNA tether held together a pair of beads (bottom pair of beads in **Fig. 5c and Supplemental Movie 2**), so we infer that the coaligned DNA stabilizes this mode of binding. Supporting this conclusion, if we crossed the DNA tethers between two pairs of beads, bridging frequently initiated at or near the crossing point (21 of 23 trials; **Fig. s9a**). Because the crossing points are expected to constrain and align the stretched tethers, the preferential initiation of bridges at these locations indicates that the DNA configuration contributes to stable protein binding.

When present, dangling DNA molecules (i.e., those that had both ends bound to only one bead) were progressively bundled together with the stretched DNA tethers until no more DNA could be coaligned (**Fig. 5c and Supplemental Movie 2**). This bundling indicates that binding of the RM-TDB domain can exert force to overcome the displacement of the dangling DNA by the flow.

The protein dissociated rapidly when preassembled bundles were moved to a protein-free channel (0.034 ± 0.014 s^-1^; **Fig. 5d,e, Fig. s9b, and Supplemental Movie 3**). Coincident with protein dissociation, coaligned tethers came apart (**Fig. 5e**). Protein binding could also be reversed by pulling the beads apart, which resulted in abrupt transitions in force-extension curves (**Fig. 5f**), or by holding the traps in fixed position with high initial tension on the tethers (**Fig. s9c**). It is plausible that, because different DNA molecules are anchored at different places on the beads, pulling the beads apart exerts tangential forces that pull the duplexes apart, leading in turn to unbundling and protein release.

We conclude that the RM-TDB domain has a bridging activity that is able to bundle DNA molecules together reversibly. We further infer that protein binding and coalignment of the DNA mutually reinforce one another to promote cooperative assembly of nucleoprotein filaments.

### DNA binding by large RM-TDB assemblies

Although we did not observe coating of single DNA duplexes by RM-TDB (**Fig. 5c**), we did frequently observe the sudden appearance of bright protein foci that then remained stably bound to their initial locations on DNA tethers (**Fig. 5g, Fig. s10a, and Supplemental Movie 4**). Each focus contained multiple copies of the RM-TDB domain, as judged by analysis of fluorescence intensity, and showed little or no indication of coinciding with a spot of locally condensed DNA (**Fig. s10b**). Moreover, fluorescence intensity was already maximal when a focus first appeared, showing little or no evidence of net growth by addition of more protein (**Fig. 5g,h and Fig. s10b,c**). When beads tethered by a protein-bound DNA molecule were pulled apart at constant velocity, multiple transitions in the force-extension curves could be detected (**Fig. s10d,e,f**), suggesting that interactions between distinct DNA segments were disrupted (**Fig. s10d,e,f**). We infer that these focal binding events reflect capture by the DNA of rare, relatively large, pre-existing RM-TDB assemblies that can bind simultaneously to multiple segments along the same DNA molecule. These assemblies may be nonspecific protein aggregates, specific multiprotein complexes, or nucleoprotein condensates formed on trace nucleic acid in the purified preparations.

More importantly, we observed a different mode of protein binding on dangling DNA molecules in which relatively modest initial protein fluorescence at the tip of the DNA increased in intensity over time, indicating incorporation of new proteins over time **(Fig. 5h,i, Fig. s10c, and Supplemental Movie 5)**. Protein-bound DNA tips moved progressively upward against flow toward the beads, coincident with the increase in signal intensity (**Fig. 5i, s10g**). Tip binding could also occur coincidentally with apparent bridging of the parallel arms of a single dangling DNA (**Fig. s10g)**.

These findings with dangling DNA suggest that binding of the RM-TDB domain can nucleate at or near positions where segments of a single DNA duplex fold back in parallel, perhaps through bridging of the coaligned stretches of DNA segments. This nucleation can then lead to progressive accumulation of more protein and incorporation of more of the unconstrained DNA. These nucleoprotein structures are able to exert force on the DNA as they assemble, as indicated by their ability to pull DNA up toward the bead against flow (**Fig. 5i**). As this mode of binding was not seen with extended DNA tethers, it suggests that the formation and growth of these protein-DNA assemblies are fostered by the availability of the less constrained dangling DNA. These binding events thus have properties expected for nucleation and growth of condensates that are similar to those that form when the protein is allowed to assemble on unconstrained DNA in solution.

### Evolutionarily variable DNA-binding surface of the RM-TDB domain

We previously showed that DNA binding by full-length RM complexes was compromised by alanine substitution of four basic residues (R395, K396, K399, and R400) within the RM-TDB domain (Claeys Bouuaert et al., 2021a). This “Rec114-4KR” mutant reduced binding to an 80-bp DNA substrate by about 1.5-fold, diminished condensate formation on plasmid substrates, attenuated formation of chromatin-bound foci *in vivo*, and almost completely eliminated DSB formation during meiosis.

Our model provides a structural framework for understanding how these residues and others contribute to DNA binding. Overall, the surface of the RM-TDB domain is highly positively charged because of symmetric faces of the Rec114_C_ dimer that display outward-directed basic side chains (**Fig. 6a**). Each face comprises the Rec114-4KR residues R395, K396, K399, and R400 plus residues K403 and K407 from one Rec114_C_ chain, along with residues K417 and K424 from the other chain (**Fig. 6a**). K395 and K396 lie in the turn between helices 1 and 2; K399, R400, K403, and K407 lie within helix 2; and K417 and K424 lie within helix 3 (**Fig. 3a and Fig. 6a**).

**Fig. 6:**
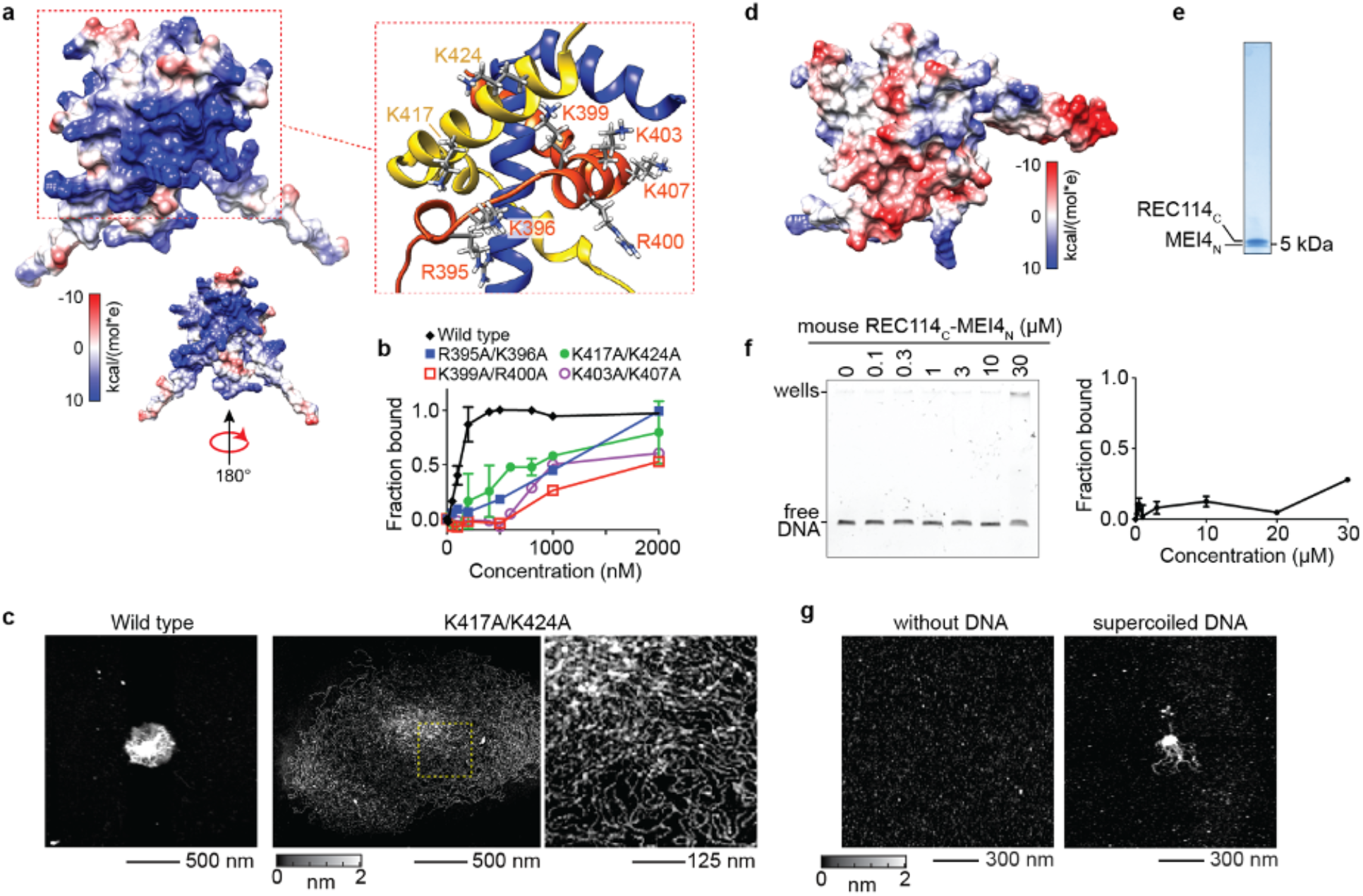
DNA-binding surface of the RM-TDB domain. (a) Electrostatic surface potential. The zoomed structural model highlights key positively charged residues. (b) EMSAs of wild type and double-alanine substitutions (complexes of Rec114_388-428_ with Mei4_N_). Error bars are mean ± range from two experiments. C_50_ values were 110 ± 20 nM (wild type); 1000 ± 50 nM (K403A/K407A); 900 ± 100 nM (K417A/K424A). Both R395A/K396A and K399A/R400A EMSAs were conducted once (C_50_ of ~1 μM and ~ 2 μM, respectively). (c) AFM images of DNA condensation by wild-type (500 nM) or K417A/424A mutant (2 μM) RM-TDB domain (complexes of Rec114_388-428_ with Mei4_N_). The 1000 bp DNA was at 1 ng/μl. The region in the dashed box is shown at higher magnification to the right. (d) Electrostatic surface potential of the mouse RM-TDB domain. (e) SDS-PAGE of purified mouse RM-TDB domain. (f) EMSA of mouse RM-TDB domain binding to a 150-bp DNA substrate. Error bars indicate mean ± range from two experiments; binding was not saturated at 30 μM. (g) AFM imaging of 6 μM mouse RM-TDB domain in the absence (left) or presence (right) of 1.7 ng/μl supercoiled pUC19 plasmid DNA.

We tested the contributions of these residues to DNA binding by the minimal RM-TDB domain using double alanine substitutions. Mutation of K399/R400 substantially reduced the apparent affinity for a 150-bp DNA substrate in EMSAs (C_50_ of ~2 μM; **Fig. 6b and Fig. s11a**). Mutation of R395/K396, K403/K407 or K417/K424 also reduced binding, but to a lesser extent (C_50_ between 500 nM and 1 μM). We also examined condensate formation by AFM for the K417A/K424A mutant. Interestingly, this mutant at 2 μM was still able to gather 1000-bp linear DNA into large clusters, but with a less compact higher order structure of larger radius and reduced height (**Fig. 6c and Fig. s11b,c**). These findings suggest that electrostatic interactions are crucial for the RM-TDB domain to assemble densely packed nucleoprotein cores.

Unlike the residues that contribute to protein-protein interactions within the folded RM-TDB core, these outward-facing basic residues are highly variable between Rec114 orthologs (**Fig. 3a**). Among the species examined here, only the *S. cerevisiae* protein is predicted to have such a strongly positive electrostatic surface potential (**Fig. 6d and Fig. s11d**).

Interestingly, the mouse RM-TDB domain is predicted to have an overall negative surface potential, with only a few basic surface residues (**Fig. 6d**). To test whether the mouse protein is able to bind and condense DNA, we expressed and purified trimeric complexes of REC114 residues 203–259 (mREC114_C_) with MEI4 residues 1–42 (mMEI4_N_) (**Fig. 6e and Fig. s11e**). This protein complex bound to a 150-bp DNA substrate and shifted it to the wells in EMSAs, but with much lower apparent affinity than the yeast protein (**Fig. 6f**). By AFM, the mouse RM-TDB protein appeared as fairly uniform, small particles in the absence of DNA, but was also able to form nucleoprotein clusters when incubated with plasmid DNA (**Fig. 6g**). We conclude that the DNA binding and condensation activity is conserved in the mouse protein, albeit with substantial differences in affinity and the details of the putative protein-DNA interface.

## Discussion

In this study, we focused on the minimal folded domain that forms the core of the trimeric Rec114–Mei4 interface. Computational modeling revealed a unique but evolutionarily conserved helical fold in which the C-terminal segments from a pair of Rec114 proteins embrace one another to form a roughly two-fold symmetric sleeve into which the first helix of a helix-turn-helix segment from Mei4 inserts, breaking the symmetry. We validated key predictions of the computational model by NMR experiments on this minimal RM-TDB domain from *S. cerevisiae*.

This structure raises the question of how the complex is assembled *in vivo*. Since Rec114 dimers can form readily without Mei4 and might compete with the correctly assembled RM complex for binding to the Spo11 core complex, a co-translational assembly mechanism might be beneficial (Shiber et al., 2018). In fact, overexpression of Rec114 alone results in dominant-negative inhibition of DSB formation (Bishop et al., 1999), emphasizing the potential importance of forming stoichiometric RM complexes.

Interfacial residues—particularly hydrophobic ones that contribute to both homotypic and heterotypic intermolecular interactions—are the principal contributors to the signatures of sequence conservation in these parts of both proteins. Our findings thus provide a fine-grained molecular framework for understanding the coevolution of these proteins, explaining in turn how they often act as a single functional unit during meiotic DSB formation (Li et al., 2006; Maleki et al., 2007; Kumar et al., 2018; Hinman et al., 2021; Vrielynck et al., 2021). Nonetheless, it is remarkable that many residues in this domain are highly variable between species.

The RM-TDB domain exhibits multiple DNA-binding modes distinguishable by experimental approach, including nucleoprotein condensate formation (**Fig. 4c,d and Fig. 5i**), bridging of coaligned DNA duplexes (**Fig. 5b,c**), and formation of either small, discrete particles (**Fig. 4g**) or large focal structures (**Fig. 5g**). As discussed below, we consider it likely that these DNA-binding modes are functionally related and reveal specific characteristics that underlie the formation of higher order DNA assemblies by the RM complex.

The minimal yeast RM-TDB domain was previously shown to bind DNA, but unexpectedly we also found that it is sufficient to form higher order nucleoprotein condensates *in vitro*. Although the full-length RM complex is much more efficient at making condensates, our findings point to the RM-TDB domain as an important basal module of condensate formation. The mouse and yeast proteins share this property, albeit with quantitatively very different abilities to support efficient condensation.

A critical factor for observing condensates appears to be the availability of long DNA substrates that are unconstrained (as in our AFM and confocal studies of structures formed in solution before immobilization on a surface) or minimally constrained (as with the dangling DNA in optical trap experiments). Confocal microscopy uncovered an interesting substructure of condensates in which a protein- and DNA-dense core is surrounded by a less dense nucleoprotein meshwork. The biophysical differences between these two zones are not currently clear, but previous experiments indicated that RM condensates can transition to a more stable, possibly gel-like state over time. Thus, it is possible that these two zones are related to differences between more sol-like vs. more gel-like condensates made by the full-length proteins (Claeys Bouuaert et al., 2021a).

In optical trap experiments, we observed progressive assembly of nucleoprotein structures on dangling DNA that involved net accumulation of protein concomitant with incorporation of additional DNA. It is plausible that these assemblies are related to the condensates we observed by AFM and confocal microscopy, possibly being equivalent to early dynamic steps in the formation of mature condensates.

The DNA bridging activity we also observed in these single-molecule experiments was particularly striking and unexpected. One defining feature of this mode of binding is that at least one of the DNA duplexes is physically constrained in an outstretched configuration. Although we lack evidence that bridging also occurs within condensates, it is straightforward to infer that it does so and is in fact an important component of the network of interactions that makes up a condensate. This is because the same bridging interactions that give rise to linear nucleoprotein filaments on stretched DNA would be expected to give a more complex three-dimensional meshwork on minimally or unconstrained DNA in solution.

Several properties of DNA bridging stand out, including its dynamic nature (readily reversible with fast off rate); its formation of long, contiguous stretches of coaligned DNA duplexes; its ability to bundle several DNA duplexes; and its ability to apply force and in turn to be disrupted by force. These properties, combined with the structure of the RM-TDB domain, point to two further implications. First, the rotational symmetry of the Rec114_C_ dimer, with its potential to provide two nearly identical DNA-binding faces, leads us to envision that the basal bridging unit may be a single RM-TDB trimer. However, we cannot exclude that a dimer of trimers or other higher order assembly is the base unit.

Second, we consider that a significant contributor to the cooperativity of DNA binding is the combination of DNA persistence length with a bivalent DNA-binding protein. Supporting this idea, we concluded above that protein binding and DNA coalignment mutually reinforce because pre-aligning the DNA favors protein binding and, conversely, binding of additional protein exerts force to align otherwise separate duplexes. This type of cooperativity could be entirely independent of contacts between adjacent bridging units (analogous to the ties on a railroad track), because presence of a bridge increases the effective local concentration of an adjacent protein binding site and reduces the entropic cost of forming an adjacent bridge (Wiggins et al., 2009). Alternatively, cooperativity may also be fostered by direct protein-protein interactions.

Protein-protein interactions might contribute to the formation of the small clusters of RM-TDB proteins on single DNA duplexes observed by AFM under sub-saturating conditions (**Fig. 4g**). These clusters are unlikely to represent bridging of folded back duplex DNA because the internal binding events showed no evidence of the consistent sharp DNA bends that would be expected for foldback events. Interestingly, however, the shorter contour length of cluster-bound DNA suggests that the DNA is condensed, possibly following a superhelical trajectory. Although the relationship between these clusters and other modes of DNA binding is unclear, we speculate that clusters may resemble the initial binding events that lead to bridging, condensation, or both.

The nature is also unclear for the remaining mode of DNA binding we observed, namely, the large RM-TDB foci that appeared suddenly on DNA tethers in optical trap experiments. These foci were maximally bright when they first appeared, bound stably to their initial binding sites, did not appear to incorporate substantial amounts of condensed DNA, and neither gained nor lost protein subunits at an appreciable rate. It thus appears plausible that they are non-specific, preformed protein aggregates, although we cannot exclude that these represent a rapid initial condensation within the imaging frame interval (which was typically a few seconds) that is limited by the available slack in the DNA tether. Regardless of how these DNA-binding events arise, however, we infer that each focus likely has multiple DNA binding interfaces because of their force-sensitive ability to connect distant DNA segments on the same DNA molecule. Thus, these foci may provide insight into how large multivalent RM assemblies could interact with DNA.

An important observation in our study is that most of these nucleoprotein structures generate force that can reel in more DNA or bundle DNA molecules together against opposing forces. Conversely (and by necessity), interactions of the RM-TDB domain with DNA are modulated by applied force. By extension, we infer that the more complex condensates containing RM complexes and other proteins *in vivo* also both respond to and impose force on chromatin. There is a growing appreciation of the importance of capillary forces imposed by biomolecular condensates (Gouveia et al., 2022), including forces on DNA from nucleoprotein assemblies such as those made by transcription factors (Quail et al., 2021; Renger et al., 2021; Nguyen et al., 2022).

During meiosis, mechanical stress on chromosomes has been proposed to regulate the number and spatial patterning of recombination events (Kleckner et al., 2004; Zhang et al., 2014). More recently, diffusion-based coarsening models have been proposed to explain the patterning of meiotic crossovers (Zhang et al., 2018; Morgan et al., 2021; Haversat et al., 2022) and—via RM and Mer2 condensates—DSBs (Claeys Bouuaert et al., 2021a). We therefore propose that the RM-TDB domain, with its intrinsic force-generating and force-responsive properties, is a fundamental building block that organizes meiotic recombination and that may reconcile mechanical stress and coarsening models for meiotic chromosome behavior.

## Materials and Methods

### Expression and purification of Rec114-Mei4 complexes

All yeast and mouse Rec114 and Mei4 constructs were cloned into a pETDuet-based expression vector with a 6XHis-SUMO tag, except for Rec114_C_ alone, which was cloned into a pSMT3-based vector (**Supplementary Table s1**). The plasmids were transformed into BL21(DE3) cells (Invitrogen) for overexpression. Purification of all constructs reported in this work followed the same procedure. Typically, cells were induced with 1 mM isopropyl β-d-1-thiogalactopyra-noside (IPTG) at an OD_600_ of ~0.6–0.8 for 3–4 hours at 37 °C. The cells were lysed by sonication in 25 mM HEPES-NaOH pH 7.4, 500 mM NaCl, 1× Complete protease inhibitor tablet (Roche) and 0.1 mM phenylmethanesulfonyl fluoride (PMSF) and centrifuged at 5,000g for 10 min. Cleared extract was loaded onto 1 ml preequilibrated Cobalt resin (Thermo Scientific). The column was washed extensively with wash buffer (25 mM HEPES-NaOH pH 7.4, 500 mM NaCl, 5 mM imidazole, 0.1 mM PMSF). The tagged complexes were then eluted in elution buffer containing 200 mM imidazole. The elution was dialyzed in 25 mM HEPES-NaOH pH 7.4, 200 mM NaCl overnight at 4 °C with ~0.1 mg/ml homemade Ulp1 to cleave off the His-SUMO tag. After dialysis, the sample was clarified and concentrated, then chromatographed on a Superdex 200 Increase 10/300 GL column (Cytiva) preequilibrated in 25 mM HEPES-NaOH pH 7.4, 100 mM NaCl. The fractions after size exclusion chromatography were checked by SDS-PAGE and pooled. Protein concentration was determined by A280. Concentrations were calculated on the basis of a 2:1 stoichiometry for trimeric complexes or on the basis of a dimer of Rec114_C_ only. Aliquots were frozen in liquid nitrogen and stored at −80 °C.

For {^15^N} or {^15^N-^13^C} labeled samples, a single colony was inoculated in 5 ml Luria-Bertani (LB) liquid medium and grown at 37 °C for 6–8 hours, then the culture was diluted 100-fold into 100 ml M9 minimal medium with ^15^N ammonium chloride or with both ^15^N ammonium chloride and ^13^C D-glucose from Cambridge Isotopes and grown overnight at 37 °C. The overnight culture was then transferred to 900 ml fresh M9 minimal medium and grown until OD_600_ of ~0.6–0.8 before IPTG induction. Remaining expression and purification procedures were the same as for unlabeled samples.

### Fluorescent labeling of the RM-TDB domain

Site-specific labeling was done as described previously (Wasserman et al., 2019). Briefly, 1 μM of RM-TDB in which Mei4_N_ has a C-terminal ybbR tag was incubated with ~3 μM Sfp and 2 μM of CoA-LD650 in buffer HM (50 mM HEPES-NaOH pH 7.4, 10 mM MgCl_2_) at room temperature for 2 hours in a total volume of 100 μl. The sample was then subjected to size exclusion chromatography to remove Sfp and unincorporated dye. Aliquots were frozen in liquid nitrogen and stored at –80 °C.

### NMR spectroscopy

Unless otherwise noted, NMR data were collected in 25 mM HEPES-NaOH pH 7.4, 100 mM NaCl, 0.5 mM EDTA, 1 mM TCEP, 0.05% NaN3 at 25 °C on a Bruker Avance III spectrometer at the New York Structural Biology Center (NYSBC). Rec114_C_–Mei4_N_ was assigned using 600 μM uniformly {^15^N-^13^C} labeled protein at 800 MHz (^1^H) using non-uniformly sampled HNCACB, CBCA(CO)NH, HNCA, HNCACO, HNCO, HN(COCA)NH, and H(CCCONH)-TOCSY. NOESY-HSQC data were collected using τ_mix_ = 100 ms. Assignments from Rec114_C_ Mei4_N_ were transferred directly to Rec114_388-428_–Mei4_N_ spectra and corroborated using 408 μM uniformly {^15^N-C} labeled protein at 25 °C at 700 MHz (^1^H) using non-uniformly sampled HNCACB, HNCA, HNCACO, HNCO, HN(COCA)NH, and H(CCCONH)-TOCSY. Rec114_388-428_–Mei4_13-43_ was assigned using 470 μM uniformly {^15^N-^13^C} labeled protein in 25 mM NaHPO_4_, pH 6.1, 100 mM NaCl, 0.5 mM EDTA, 1 mM TCEP, 0.05% NaN_3_, 5% D_2_O at 800 MHz (^1^H) using non-uniformly sampled HNCACB, CBCA(CO)NH, HNCA, HNCACO, HNCO, and HN(COCA)NH. Spectra for isolated Rec114_375-428_ were collected using 100 μM protein at 500 MHz (^1^H) (Weill Cornell NMR Core). All spectra were processed using NMRpipe (Delaglio et al., 1995) and reconstructed using SMILE-NMR (Ying et al., 2017) on NMRbox (Maciejewski et al., 2017). Data were analyzed using NMRFAM-sparky (Lee et al., 2015). TALOS-N was used to generate secondary structure predictions (Shen and Bax, 2013).

### CD spectroscopy

CD spectra were collected using an AVIV Biomedical Model 410 CD Spectrometer. Spectra were collected at 25 °C using one nm wavelength steps going from 300 nm to 190 nm. Each spectrum was collected using a two-minute temperature equilibration and one scan for each step with a 5 second averaging time using 0.2 mm path length plates. The concentrations for Rec114_C_-Mei4_N_ and Rec114_C_ alone were 30 μM and 100 μM, respectively.

### Bioinformatic analysis

We used IUPRED3 (Erdos et al., 2021) and ANCHOR (Dosztányi et al., 2009) to predict protein disorder. Sequence-based secondary structure prediction was done via PSIPRED 4.0 (Buchan and Jones, 2019). The multiple sequence alignment (MSA) was performed using the MAFFT 7 online server with FFT-NS-2 option (Katoh et al., 2019). The sequence and secondary structural conservation rendering figure was generated by Espript 3.0 (Robert and Gouet, 2014).

The structural models were generated by ColabFold (Mirdita et al., 2022) with a python Jupyter notebook:

https://colab.research.google.com/github/sokrypton/ColabFold/blob/main/beta/AlphaFold2_advanced.ipynb MSA was generated using mmseqs2 (Steinegger 2017). The default pair MSA and filter options were used to generate five models with the highest pLDDT scores. These models are highly similar to each other and the one with the highest score was selected for subsequent analyses.

### DNA substrates and EMSAs

Short linear DNA substrates were generated by annealing complementary oligonucleotides (sequences listed in **Supplementary Table s2**). The substrates were the following (with oligo names in parentheses): dsDNA20 (KL020 and KL021), dsDNA80 (KL024 and KL025). Oligos were mixed in equimolar concentrations (10 μM) in STE (100 mM NaCl, 10 mM Tris-HCl pH 8, 1 mM EDTA), heated and slowly cooled on a PCR thermocycler (98 °C for 3 min, 75 °C for 1 h, 65 °C for 1 h, 37 °C for 30 min, 25 °C for 10 min).

Larger linear substrates were prepared by PCR amplification of a lambda DNA template (New England Biology). Substrates were as follows: 150 bp (KL001 and KL003), 1,000 bp (KL001 and KL004). Fluorescently labelled substrates were prepared by PCR amplification of lambda as follows: Cy3–1000 bp (KL001 and KL010). PCR products were purified by agarose gel electrophoresis.

Binding reactions (10 μl) were carried out in 25 mM Tris-HCl pH 7.4, 100 mM NaCl. Unless stated otherwise, reactions contained 5 ng substrate and the indicated concentration of protein. Complexes were assembled for 20 min at room temperature and separated on 6% DNA retardation gels (Thermo Fisher scientific) at 200 V for 1–2 h. Gels were stained with SYBR safe (Invitrogen) and scanned using a ChemiDoc Imaging System (Bio-Rad). We quantified apparent affinities from protein titration EMSA experiments by linear interpolation as the concentration of protein at which 50% of the substrate was bound (referred to as the C_50_). Because we do not know the stoichiometry of protein-bound DNA or the number of protein binding sites per DNA molecule, these are only rough approximations of true Kd values and are used to provide a means of comparison between different DNA binding experiments.

### AFM imaging

For AFM imaging, protein complexes were diluted to the indicated concentration (70 nM to 6 μM) in the presence of ~1 ng/μl of different DNA substrates in 25 mM HEPES-NaOH pH 7.4, 5 mM MgCl_2_, 50 mM NaCl, 10% glycerol. Complexes were assembled at room temperature for 30 min. A volume of 40 μl of the protein–DNA binding reaction was deposited onto freshly cleaved mica (SP1) for 2 min. The sample was rinsed with 10 ml ultrapure deionized water and the surface was dried using a stream of nitrogen. AFM images were captured using an Asylum Research MFP-3D-BIO (Oxford Instruments) microscope in tapping mode at room temperature. An Olympus AC240TS-R3 AFM probe with resonance frequencies of approximately 70 kHz and spring constant of approximately 1.7 N/m was used for imaging. Images were collected at a speed of 0.5–1 Hz with a resolution of ~1 nm/pixel.

For the experiments at low protein concentration (70 nM), we used a MATLAB (2021b) script to trace the DNA and determine the contour length. Briefly, the DNA molecule boundaries were determined by the bwboundaries.m function. The contour of each DNA molecule was further determined by the bwskel.m function to measure the contour length. For complex DNA molecules with multiple RM binding sites, the bwmorph.m function was used to determine the branch points and different binding segments. Contour length was then measured for each segment. All contour length measurements were manually checked by ImageJ (Schneider et al., 2012).

For volume measurement, boundaries (B) were first determined for each DNA molecule as described above. The volume was then defined by: *V* = *rx* * *ry* * (∑_*p∈B*_(*h_p_* – *h_b_*)), where *rx* and *ry* are the resolution of each pixel, *h_p_* is the height of each pixel within the boundaries B, and *h_b_* is the average height of the background.

To estimate the number of RM-TDB proteins bound, an average protein density of 1.44 g/cm^3^ was used (Fischer et al., 2009; Bouuaert et al., 2020). For a single RM-TDB domain, the volume Vp was estimated to be ~20.7 nm^3^ and there was an average of 2.2 particles per DNA molecule (N_p_ in **Fig. 4h**). The average volume difference (V_d_) between in the absence and presence of RM-TDB was 303 nm^3^, so the average number of RM-TDB bound was estimated as 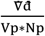, which is 6.7.

### Scanning confocal microscopy

Reactions (20 μl) of 450 nM Rec114_C_–Mei4_N_-LD650 mixed with 25 nM Cy3-1000 bp linear DNA in 25 mM HEPES-NaOH pH 7.4, 100 mM NaCl were incubated at room temperature for 10 min and deposited onto the surface of a coverslip. The confocal image data were collected with a LUMICKS C-trap instrument using 532 nm and 638 nm lasers to excite the Cy3 and LD650 dyes, respectively. Two-dimensional scan movies were recorded by BlueLake software (LUMICKS) at roughly ~200 μs/pixel and 100 nm/pixel resolution.

Confocal images were split into LD650 (protein) and Cy3 (DNA) channels and analyzed separately. The sizes of the whole condensates and of just the dense core were determined by ImageJ using thresholding. The radius of the core was then determined simply by *A* = *πr*^2^, where A is the area of the core and r is the radius. The core radii were similar in the two channels, so only the protein channel data were reported.

To determine the thickness of the halo structure, we used a custom MATLAB script. Briefly, the boundaries of the cores and the whole condensates were determined by thresholding. The center of the core was also determined. A series of lines at 10° angles were then drawn starting from the center point. The halo thickness along each line was determined and then averaged for a given condensate to generate each data point in **Fig. 4f.**

### Single-molecule optical trap experiments: data acquisition

Single-molecule experiments were conducted at room temperature on a LUMICKS C-Trap instrument combining three-color confocal fluorescence microscopy with quadruple-trap optical tweezers. Laminar flow separated channels 1–3, which were used to form DNA tethers between 4.89-μm streptavidin-coated polystyrene beads (Spherotech) held in traps with a stiffness of ~0.3 pN/nm. Under constant flow, a single bead was caught in each trap in channel 1. The traps were then quickly moved to channel 2 containing the biotinylated lambda DNA (LUMICKS). By moving one trap against the direction of flow but toward the other trap, and vice versa, a DNA tether could be formed and detected via a change in the force-distance (FD) curve. The traps were then moved to channel 3 containing only PBS buffer for force calibration without flow, and the presence of a single DNA was verified by the FD curve. Orthogonal channels 4 and 5 served as protein loading and/or experimental imaging chambers as described for each assay. The flow was turned on to visualize binding of the RM-TDB domain. SYTOX Orange and LD650 were excited by two laser lines at 532 nm and 638 nm, respectively. The imaging buffer condition was 25 mM HEPES-NaOH, pH 7.4, 50 mM NaCl, 5 mM MgCl_2_, 10% glycerol and 5 nM SYTOX Orange if not specified otherwise. The RM-TDB domain concentration in all experiments shown here was 20 nM.

For experiments with multiple DNA tethers and/or dangling DNA, extra waiting time was spent in channel 2 to allow more biotinylated lambda DNA molecules to bind to the bead surface. To generate crossed DNA, a LUMICKS Q-trap system was utilized. Briefly, four individual 4.89-μm streptavidin-coated polystyrene beads were trapped by four laser traps in channel 1. The trap stiffness was kept the same as above. The traps were then moved to channel 2 to catch biotinylated lambda DNA. The first pair of beads were trapped by Trap 1 and Trap 2 to catch the first DNA tether and left > 10 μm apart to prevent more tethers forming. The second pair of beads were trapped by Trap 3 and Trap 4 to catch the second tether. The traps were then moved to channel 3 to verify the DNA. To achieve single DNA tether, the DNA was held at ~60 pN for some period of time until only single tether was left, which was characterized by the FD curve. A script was used to generate the crossed DNA configuration at channel 3. The data were recorded by BlueLake software (LUMICKS).

### Single-molecule experiments: data analysis

The C-trap fluorescence data were processed and visualized by ImageJ and custom python scripts based on Pylake package (v0.13.0) provided by LUMICKS. The line tracking was performed using the track_greedy function from the Pylake package. Waiting time was extracted manually based on the starting time point of the binding from two-dimensional scan movies. The fluorescence images were visualized and exported by ImageJ.

For extraction of mean photon counts, raw data of two-dimensional scans were exported by Pylake as .tiff files and analyzed in MATLAB. A region of interest (ROI) box was drawn manually to include extract photon counts of the region, and mean photon counts of the ROI were determined by averaging the photon counts per pixel. The same size boxes were applied to the same movie. The normalized mean intensity is ratio of the mean photon count per pixel for each frame to the mean photon count per pixel for the first frame (t = 0). Dissociation rates were determined by fitting the mean photon counts to a single exponential curve.

## Code Availability

Custom MATLAB scripts are available online at: https://github.com/kliu39/RM-paper.

## Author Contributions and Notes

KL, EMG, DE, SL and SK designed research; KL, EMG and SP performed research; and KL, EMG, DE and SK wrote the paper with input from SL. The authors declare no conflict of interest.

## Acknowledgments

We thank Michael Wasserman and Ling Wang for reagents; Gabriella Chua for technical support for the confocal microscopy experiments; Biran Wang and Young Hun Kim for AFM experiments; Shibani Bhattacharyya for support in performing NMR experiments at the NYSBC; Clay Bracken for support in performing NMR experiments at the Weill Cornell NMR Core Facility; and Murray Tipping for discussion of C-trap experiments. We thank members of the Keeney and S. Liu laboratories for discussions. We thank the Memorial Sloan Kettering (MSK) Molecular Cytology core facility for assistance with AFM and C-trap experiments. MSK core facilities are supported by National Cancer Institute Cancer Center support grant P30 CA08748. D.E. is a member of the NYSBC, which is supported by ORIP/NIH facility improvement grant CO6RR015495. The 700 MHz spectrometer was purchased with funds from NIH grant S10OD018509. Data collected using the 800 MHz Avance III spectrometer are supported by NIH grant S10OD016432. Some of the work presented here was conducted at the Center on Macromolecular Dynamics by NMR Spectroscopy located at the NYSBC, supported by NIH grant GM118302. The Weill Cornell NMR Core Facility is supported by NIH grant S10 OD016320. K.L. is a Damon Runyon Fellow supported in part by the Damon Runyon Cancer Research Foundation (DRG-[2389-20]). This work was supported by NIH grants R35 GM136686 (to D.E.), DP2 HG010510 (to S.L.), R35 GM118092 (to S.K.), and R01 HD110120 (to S.K. and D. Patel).

## References

Arora, C., Kee, K., Maleki, S., and Keeney, S. (2004). Antiviral Protein Ski8 Is a Direct Partner of Spo11 in Meiotic DNA Break Formation, Independent of Its Cytoplasmic Role in RNA Metabolism. Mol. Cell 13, 549–559.

Bishop, D.K., Nikolski, Y., Oshiro, J., Chon, J., Shinohara, M., and Chen, X. (1999). High copy number suppression of the meiotic arrest caused by a *dmc1* mutation: *REC114* imposes an early recombination block and RAD54 promotes a *DMC1* -independent DSB repair pathway. Genes to Cells 4, 425–444.

Boekhout, M., Karasu, M.E., Wang, J., Acquaviva, L., Pratto, F., Brick, K., Eng, D.Y., Xu, J., Camerini-Otero, R.D., Patel, D.J., et al. (2019). REC114 Partner ANKRD31 Controls Number, Timing, and Location of Meiotic DNA Breaks. Mol. Cell 74, 1053–1068.e8.

Buchan, D.W.A., and Jones, D.T. (2019). The PSIPRED Protein Analysis Workbench: 20 years on. Nucleic Acids Res. 47, W402–W407.

Carballo, J.A., Panizza, S., Serrentino, M.E., Johnson, A.L., Geymonat, M., Borde, V., Klein, F., and Cha, R.S. (2013). Budding Yeast ATM/ATR Control Meiotic Double-Strand Break (DSB) Levels by Down-Regulating Rec114, an Essential Component of the DSB-machinery. PLoS Genet. 9.

Claeys Bouuaert, C., Pu, S., Wang, J., Oger, C., Daccache, D., Xie, W., Patel, D.J., and Keeney, S. (2021a). DNA-driven condensation assembles the meiotic DNA break machinery. Nature 592, 144–149.

Claeys Bouuaert, C., Tischfield, S.E., Pu, S., Mimitou, E.P., Arias-Palomo, E., Berger, J.M., and Keeney, S. (2021b). Structural and functional characterization of the Spo11 core complex. Nat. Struct. Mol. Biol. 28, 92–102.

Delaglio, F., Grzesiek, S., Vuister, G., Zhu, G., Pfeifer, J., and Bax, A. (1995). NMRPipe: A multidimensional spectral processing system based on UNIX pipes. J. Biomol. NMR 6, 277–293.

Dosztányi, Z., Mészáros, B., and Simon, I. (2009). ANCHOR: Web server for predicting protein binding regions in disordered proteins. Bioinformatics 25, 2745–2746.

Erdos, G., Pajkos, M., and Dosztányi, Z. (2021). IUPred3: Prediction of protein disorder enhanced with unambiguous experimental annotation and visualization of evolutionary conservation. Nucleic Acids Res. 49, W297–W303.

Fischer, H., Polikarpov, I., and Craievich, A.F. (2009). Average protein density is a molecular-weight-dependent function. Protein Sci. 13, 2825–2828.

Gouveia, B., Kim, Y., Shaevitz, J.W., Petry, S., Stone, H.A., and Brangwynne, C.P. (2022). Capillary forces generated by biomolecular condensates. Nature 609, 255–264.

Greenfield, N.J. (2007). Using circular dichroism spectra to estimate protein secondary structure. Nat. Protoc. 1, 2876–2890.

Guo, H., Stamper, E.L., Sato-Carlton, A., Shimazoe, M.A., Li, X., Zhang, L., Stevens, L., Tam, K.J., Dernburg, A.F., and Carlton, P.M. (2022). Phosphoregulation of DSB-1 mediates control of meiotic double-strand break activity. Elife 11.

Haversat, J., Woglar, A., Klatt, K., Akerib, C.C., Roberts, V., Chen, S.-Y., Arur, S., Villeneuve, A.M., and Kim, Y. (2022). Robust designation of meiotic crossover sites by CDK-2 through phosphorylation of the MutSγ complex. Proc. Natl. Acad. Sci. 119.

Henderson, K.A., Kee, K., Maleki, S., Santini, P.A., and Keeney, S. (2006). Cyclin-Dependent Kinase Directly Regulates Initiation of Meiotic Recombination. Cell 125, 1321–1332.

Hinman, A.W., Yeh, H.-Y., Roelens, B., Yamaya, K., Woglar, A., Bourbon, H.-M.G., Chi, P., and Villeneuve, A.M. (2021). Caenorhabditis elegans DSB-3 reveals conservation and divergence among protein complexes promoting meiotic double-strand breaks. Proc. Natl. Acad. Sci. 118, e2109306118.

Holm, L., and Laakso, L.M. (2016). Dali server update. Nucleic Acids Res. 44, W351–W355.

Jumper, J., Evans, R., Pritzel, A., Green, T., Figurnov, M., Ronneberger, O., Tunyasuvunakool, K., Bates, R., Žídek, A., Potapenko, A., et al. (2021). Highly accurate protein structure prediction with AlphaFold. Nature 596, 583–589.

Katoh, K., Rozewicki, J., and Yamada, K.D. (2019). MAFFT online service: multiple sequence alignment, interactive sequence choice and visualization. Brief. Bioinform. 20, 1160–1166.

Keeney, S. (2008). Spo11 and the formation of DNA double-strand breaks in meiosis. Genome Dyn. Stab. 2, 81–123.

Kleckner, N., Zickler, D., Jones, G.H., Dekker, J., Padmore, R., Henle, J., and Hutchinson, J. (2004). A mechanical basis for chromosome function. Proc. Natl. Acad. Sci. U. S. A. 101, 12592–12597.

Kumar, R., Bourbon, H.M., and De Massy, B. (2010). Functional conservation of Mei4 for meiotic DNA double-strand break formation from yeasts to mice. Genes Dev. 24, 1266–1280.

Kumar, R., Oliver, C., Brun, C., Juarez-Martinez, A.B., Tarabay, Y., Kadlec, J., and de Massy, B. (2018). Mouse REC114 is essential for meiotic DNA double-strand break formation and forms a complex with MEI4. Life Sci. Alliance 1, e201800259.

Lee, W., Tonelli, M., and Markley, J.L. (2015). NMRFAM-SPARKY: enhanced software for biomolecular NMR spectroscopy. Bioinformatics 31, 1325–1327.

Leicher, R., Osunsade, A., Chua, G.N.L., Faulkner, S.C., Latham, A.P., Watters, J.W., Nguyen, T., Beckwitt, E.C., Christodoulou-Rubalcava, S., Young, P.G., et al. (2022). Single-stranded nucleic acid binding and coacervation by linker histone H1. Nat. Struct. Mol. Biol. 29, 463–471.

Li, J., Hooker, G.W., and Roeder, G.S. (2006). Saccharomyces cerevisiae Mer2, Mei4 and Rec114 form a complex required for meiotic double-strand break formation. Genetics 173, 1969–1981.

Maciejewski, M.W., Schuyler, A.D., Gryk, M.R., Moraru, I.I., Romero, P.R., Ulrich, E.L., Eghbalnia, H.R., Livny, M., Delaglio, F., and Hoch, J.C. (2017). NMRbox: A resource for biomolecular NMR computation. Biophys. J. 112, 1529–1534.

Maleki, S., Neale, M.J., Arora, C., Henderson, K.A., and Keeney, S. (2007). Interactions between Mei4, Rec114, and other proteins required for meiotic DNA double-strand break formation in Saccharomyces cerevisiae. Chromosoma 116, 471–486.

Matthew, J.B., and Richards, F.M. (1983). The pH dependence of hydrogen exchange in proteins. J. Biol. Chem. 258, 3039–3044.

Mirdita, M., Schütze, K., Moriwaki, Y., Heo, L., Ovchinnikov, S., and Steinegger, M. (2022). ColabFold: making protein folding accessible to all. Nat. Methods 19, 679–682.

Morgan, C., Fozard, J.A., Hartley, M., Henderson, I.R., Bomblies, K., and Howard, M. (2021). Diffusion-mediated HEI10 coarsening can explain meiotic crossover positioning in Arabidopsis. Nat. Commun. 12, 4674.

Murakami, H., and Keeney, S. (2014). Temporospatial coordination of meiotic dna replication and recombination via DDK recruitment to replisomes. Cell 158, 861–873.

Nguyen, T., Li, S., Chang, J.T.-H., Watters, J.W., Ng, H., Osunsade, A., David, Y., and Liu, S. (2022). Chromatin sequesters pioneer transcription factor Sox2 from exerting force on DNA. Nat. Commun. 13, 3988.

Nore, A., Juarez-Martinez, A.B., Clément, J., Brun, C., Diagouraga, B., Laroussi, H., Grey, C., Bourbon, H.M., Kadlec, J., Robert, T., et al. (2022). TOPOVIBL-REC114 interaction regulates meiotic DNA double-strand breaks. Nat. Commun. 13, 7048.

Panizza, S., Mendoza, M.A., Berlinger, M., Huang, L., Nicolas, A., Shirahige, K., and Klein, F. (2011). Spo11-Accessory Proteins Link Double-Strand Break Sites to the Chromosome Axis in Early Meiotic Recombination. Cell 146, 372–383.

Papanikos, F., Clément, J.A.J., Testa, E., Ravindranathan, R., Grey, C., Dereli, I., Bondarieva, A., Valerio-Cabrera, S., Stanzione, M., Schleiffer, A., et al. (2019). Mouse ANKRD31 Regulates Spatiotemporal Patterning of Meiotic Recombination Initiation and Ensures Recombination between X and Y Sex Chromosomes. Mol. Cell 74, 1069–1085.e11.

Quail, T., Golfier, S., Elsner, M., Ishihara, K., Murugesan, V., Renger, R., Jülicher, F., and Brugués, J. (2021). Force generation by protein–DNA co-condensation. Nat. Phys. 17, 1007–1012.

Renger, R., Morin, J.A., Lemaitre, R., Ruer-Gruss, M., Jülicher, F., Hermann, A., and Grill, S.W. (2022). Co-condensation of proteins with single-and double-stranded DNA. Proc. Natl. Acad. Sci. 119.

Robert, X., and Gouet, P. (2014). Deciphering key features in protein structures with the new ENDscript server. Nucleic Acids Res. 42, W320–W324.

Robert, T., Vrielynck, N., Mézard, C., de Massy, B., and Grelon, M. (2016). A new light on the meiotic DSB catalytic complex. Semin. Cell Dev. Biol. 54, 165–176.

Rosu, S., Zawadzki, K.A., Stamper, E.L., Libuda, D.E., Reese, A.L., Dernburg, A.F., and Villeneuve, A.M. (2013). The C. elegans DSB-2 Protein Reveals a Regulatory Network that Controls Competence for Meiotic DSB Formation and Promotes Crossover Assurance. PLoS Genet. 9, e1003674.

Schneider, C.A., Rasband, W.S., and Eliceiri, K.W. (2012). NIH Image to ImageJ: 25 years of image analysis. Nat. Methods 9, 671–675.

Shen, Y., and Bax, A. (2013). Protein backbone and sidechain torsion angles predicted from NMR chemical shifts using artificial neural networks. J. Biomol. NMR 56, 227–241.

Shiber, A., Döring, K., Friedrich, U., Klann, K., Merker, D., Zedan, M., Tippmann, F., Kramer, G., and Bukau, B. (2018). Cotranslational assembly of protein complexes in eukaryotes revealed by ribosome profiling. Nature 561, 268–272.

Stamper, E.L., Rodenbusch, S.E., Rosu, S., Ahringer, J., Villeneuve, A.M., and Dernburg, A.F. (2013). Identification of DSB-1, a Protein Required for Initiation of Meiotic Recombination in Caenorhabditis elegans, Illuminates a Crossover Assurance Checkpoint. PLoS Genet. 9, 1–18.

Tessé, S., Bourbon, H.M., Debuchy, R., Budin, K., Dubois, E., Liangran, Z., Antoine, R., Piolot, T., Kleckner, N., Zickler, D., et al. (2017). Asy2/Mer2: An evolutionarily conserved mediator of meiotic recombination, pairing, and global chromosome compaction. Genes Dev. 31, 1880–1893.

Vrielynck, N., Schneider, K., Rodriguez, M., Sims, J., Chambon, A., Hurel, A., De Muyt, A., Ronceret, A., Krsicka, O., Mézard, C., et al. (2021). Conservation and divergence of meiotic DNA double strand break forming mechanisms in Arabidopsis thaliana. Nucleic Acids Res. 1–15.

Wasserman, M.R., Schauer, G.D., O’Donnell, M.E., and Liu, S. (2019). Replication Fork Activation Is Enabled by a Single-Stranded DNA Gate in CMG Helicase. Cell 178, 600–611.e16.

Wiggins, P.A., Dame, R.T., Noom, M.C., and Wuite, G.J.L. (2009). Protein-mediated molecular bridging: A key mechanism in biopolymer organization. Biophys. J. 97, 1997–2003.

Yadav, V.K., and Claeys Bouuaert, C. (2021). Mechanism and Control of Meiotic DNA Double-Strand Break Formation in S. cerevisiae. Front. Cell Dev. Biol. 9, 1–20.

Yin, J., Lin, A.J., Golan, D.E., and Walsh, C.T. (2006). Site-specific protein labeling by Sfp phosphopantetheinyl transferase. Nat. Protoc. 1, 280–285.

Ying, J., Delaglio, F., Torchia, D.A., and Bax, A. (2017). Sparse multidimensional iterative lineshape-enhanced (SMILE) reconstruction of both non-uniformly sampled and conventional NMR data. J. Biomol. NMR 68, 101–118.

Zhang, L., Liang, Z., Hutchinson, J., and Kleckner, N. (2014). Crossover Patterning by the Beam-Film Model: Analysis and Implications. PLoS Genet. 10, e1004042.

Zhang, L., Köhler, S., Rillo-Bohn, R., and Dernburg, A.F. (2018). A compartmentalized signaling network mediates crossover control in meiosis. Elife 7.

