## Supplementary Tables and Figures for "Structure and DNA bridging activity of the essential Rec114–Mei4 trimer interface"

**This pdf contains:**

Supplementary Tables 1 and 2.
Supplementary Figures 1– 11

**Additional information, in separate .avi files:**

Supplementary Movies 1–5

**Supplemental Table S1. Plasmids**

| Name | Description |
|---|---|
| pKL067 | HisSUMO-Rec114(375-428):Mei4(1-43) in pETDuet1 |
| pKL020 | HisSUMO-Rec114(375-428):Mei4(1-43)-ybbR in pETDuet1 |
| pSP63 | HisSUMO-Rec114(325-428):Mei4(1-90) in pETDuet1 |
| pKL080 | Rec114(386-428):HisSUMO-Mei4(1-43) in pETDuet1 |
| pKL086 | Rec114(388-428):HisSUMO-Mei4(1-43) in pETDuet1 |
| pKL077 | Rec114(399-428):HisSUMO-Mei4(1-43) in pETDuet1 |
| pKL088 | Rec114(388-428):HisSUMO-Mei4(13-43) in pETDuet1 |
| pKL110 | HisSUMO-Rec114(375-428) in pSMT3 |
| pKL075 | mREC114(203-259):HisSUMO-mMEI4(1-42) in pETDuet1 |
| pKL091 | Rec114(388-428):HisSUMO-Mei4(1-43) Rec114 R394A/K395A in pETDuet1 |
| pKL092 | Rec114(388-428):HisSUMO-Mei4(1-43) Rec114 K399A/R400A in pETDuet1 |
| pKL095 | Rec114(388-428):HisSUMO-Mei4(1-43) Rec114 K417A/K424A in pETDuet1 |
| pKL097 | Rec114(388-428):HisSUMO-Mei4(1-43) Rec114 K403A/K407A in pETDuet1 |

**Supplemental Table S2. Oligonucleotides**

| Name | Sequence |
|---|---|
| KL020 | AAAGATGTCCTAGCAATGTA |
| KL021 | TACATTGCTAGGACATCTTT |
| KL024 | CTAGTATAGAGCCGGCGCGCCATGTCTAGATAGCGTTAGGTCTGCCGAATAGTAC TACTCGGATCCCGAGCGAACCACGC |
| KL025 | GCGTGGTTCGCTCGGGATCCGAGTAGTACTATTCGGCAGACCTAACGCTATCTAG ACATGGCGCGCCGGCTCTATACTAG |
| KL010 | /5Cy3/ATTTCCACACCCTGTTTCTCCAGCGCAGCACCGTAAT |
| KL001 | ATATCGCTGCCGGGCTGGGTGT |
| KL003 | ATCTGGCTCGCCTGACGGGATGC |
| KL004 | ATTTCCACACCCTGTTTCTCCAGCGCAGCACCGTAAT |

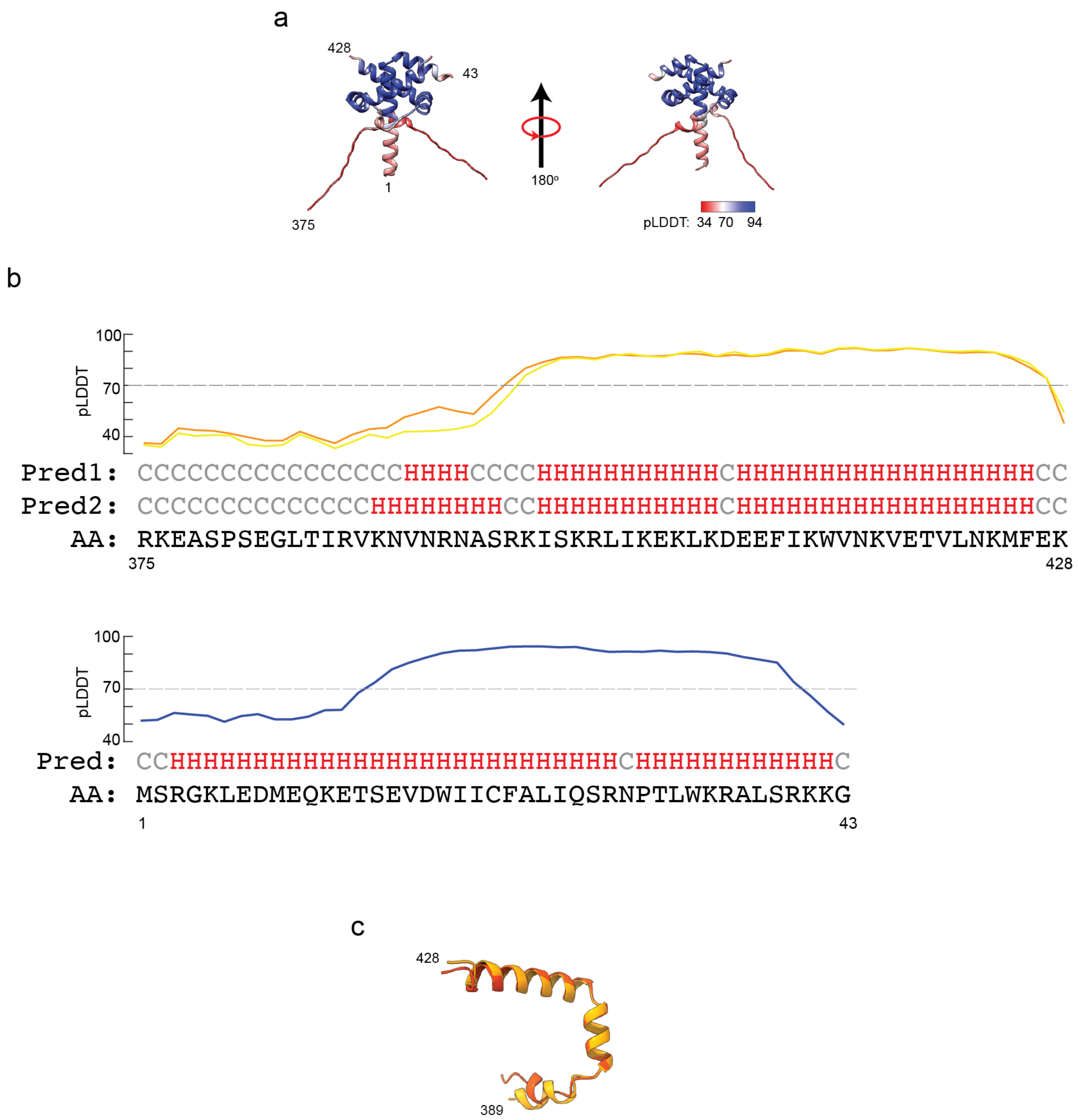


**Fig. s1: AlphaFold2 model. (a)** The AlphaFold2 model from **Fig. 1b** is color-coded by predicted local distance difference test (pLDDT) score. Residues with scores below 70 are considered low confidence. (**b**) Secondary structure prediction (Pred) and pLDDT score of the two Rec114$_C$ chains (top) and Mei4$_N$ (bottom) from the AlphaFold2 model. H: helix; C: coil. **(c)** Superimposition of $\alpha$-helices 1–3 from the two copies of Rec114$_C$ in the AlphaFold2 model.

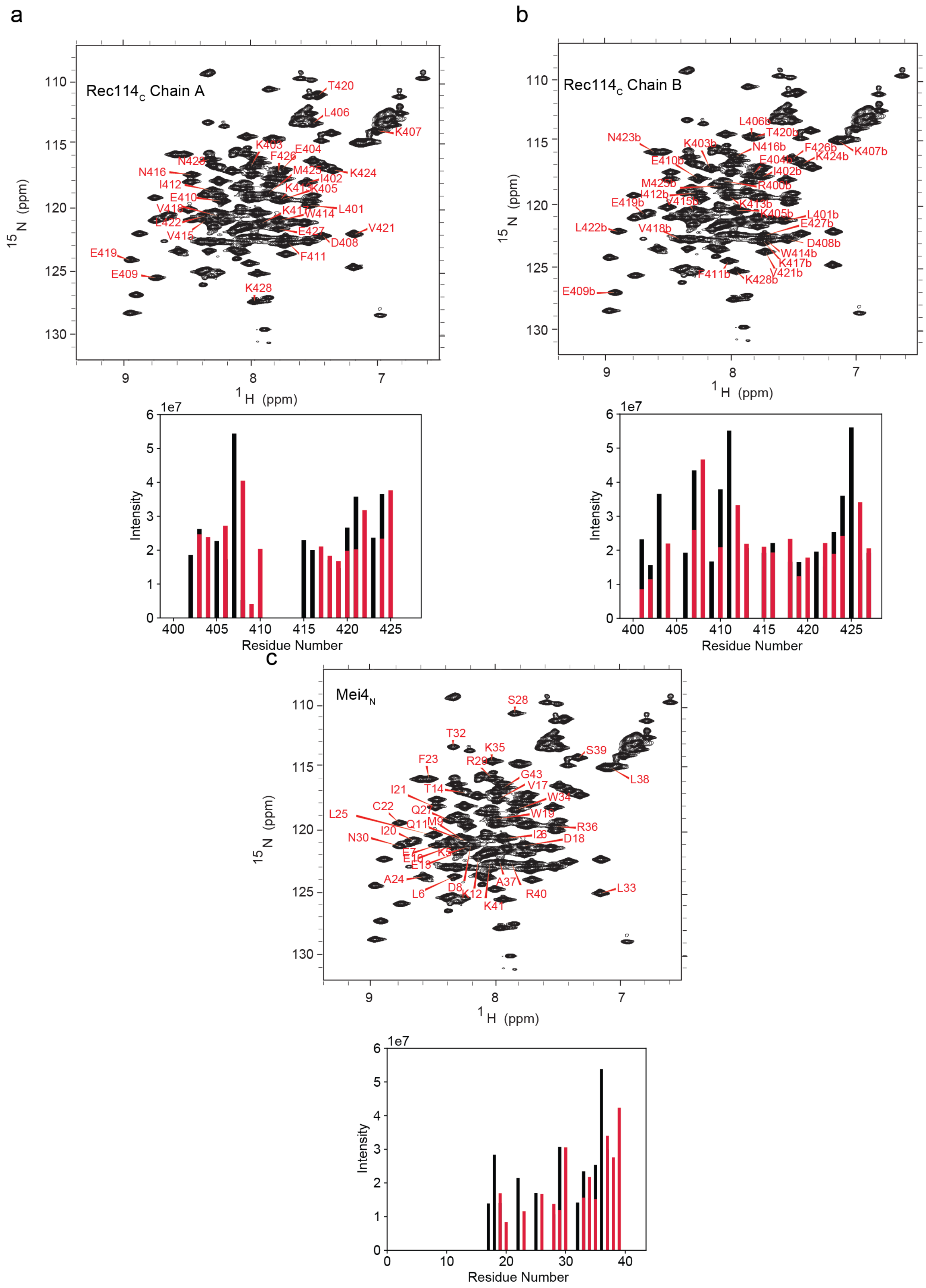
a
Rec114c Chain A
T420
L406
K407
K403
E404
F426
N428
N416
M425
K424
I402
I412
K413
K405
E410
V418
K417
W414
L401
E422
E427
V421
V415
D408
E419
F411
E409
K428
15N (ppm)
1H (ppm)
Intensity
Residue Number
1e7
b
Rec114c Chain B
L406b
T420b
N423b
K403b
N416b
F426b
K407b
E410b
E404b
K424b
I402b
M425b
R400b
I412b
E419b
V415b
K413b
K405b
L401b
E427b
L422b
V418b
D408b
W414b
K417b
F411b
V421b
K428b
E409b
15N (ppm)
1H (ppm)
Intensity
Residue Number
1e7
c
Mei4N
S28
T32
K35
S39
F23
R29
G43
L38
I21
T14
V17
W34
C22
Q27
W19
L25
Q11
M9
R36
I20
I26
D18
N30
E7
E16
K5
E13
A24
A37
L6
D8
K12
R40
L33
K41
15N (ppm)
1H (ppm)
Intensity
Residue Number
1e7

**Fig. s2: Resonance assignments and NOEs for trimers of Rec114$_C$ plus Mei4$_N$. (a, b)** Top, assigned amide resonances for Rec114$_C$ chains A and B, respectively. Note: we do not know which chain corresponds to which in the AlphaFold2 model. Bottom, i+1 (black) and i-1 (red) NH-NH NOEs for the assigned amides (residues 400–428) in each Rec114$_C$ chain. **(c)** Top, assigned chemical shifts for Mei4$_N$. Bottom, NH-NH NOEs for the amides of residues 17–40 in Mei4$_N$.

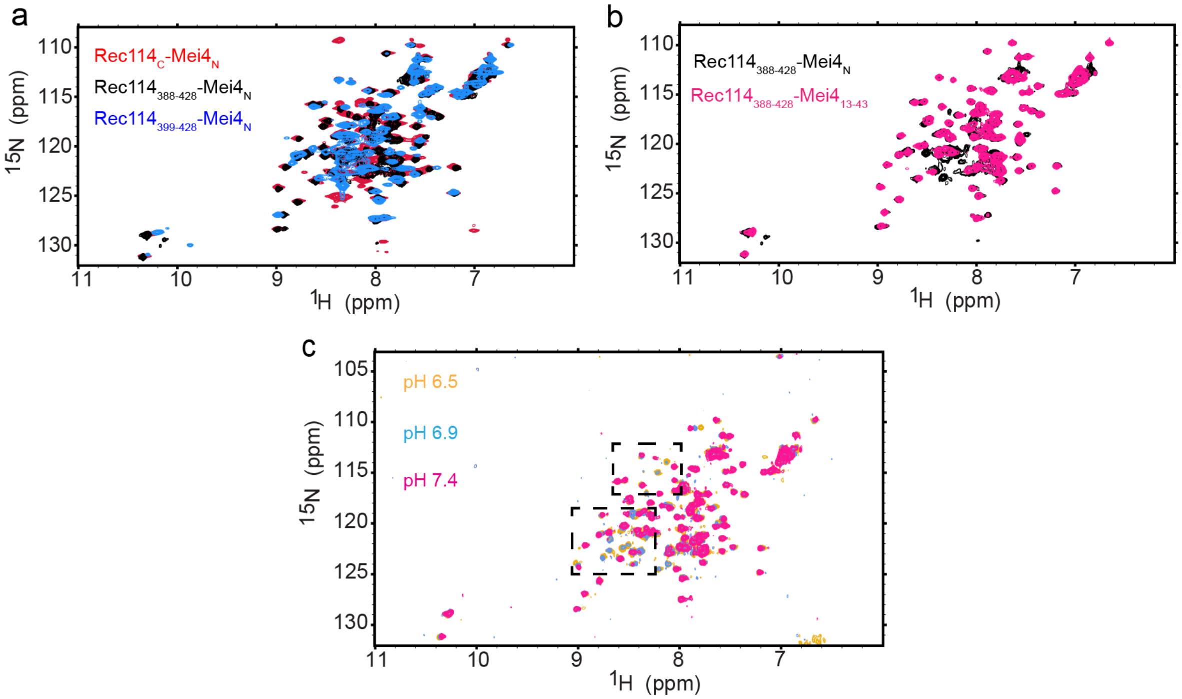


**Fig. s3: NMR studies of truncated Rec114$_C$–Mei4$_N$ complexes at varying pH values. (a)** Two-dimensional {$^1$H-$^{15}$N} HSQC spectra of complexes of Mei4$_N$ with Rec114 fragments Rec114$_C$ (red, 600 μM uniformly $^{15}$N$^{13}$C-labeled protein collected at 800 MHz ($^1$H)), Rec114$_{388\text{-}428}$ (black, 45 μM uniformly $^{15}$N-labeled protein collected at 500 MHz ($^1$H, Weill Cornell Medicine)), and Rec114$_{399\text{-}428}$ (blue, 500 μM uniformly $^{15}$N$^{13}$C-labeled protein collected at 800 MHz ($^1$H)). Removing residues 375–387 resulted in minimal changes other than the elimination of peaks corresponding to the removed residues. In contrast, removing residues 375–398 resulted in significant chemical shift perturbations and line broadening, suggestive of partial unfolding and indicating that residues 388–398 are important for the stability of the complex. **(b)** Two-dimensional {$^1$H-$^{15}$N} HSQC spectra of complexes of Rec114$_{388\text{-}428}$ with Mei4$_N$ (black, 45 μM uniformly $^{15}$N-labeled protein collected at 500 MHz ($^1$H, Weill Cornell Medicine)) or Mei4$_{13\text{-}43}$ (pink, 35 μM uniformly $^{15}$N-labeled protein collected at 500 MHz ($^1$H, Weill Cornell Medicine)). Minimal differences were observed other than elimination of signals from the removed residues. **(c)** Two-dimensional {$^1$H-$^{15}$N} HSQC spectra of 30 μM of Rec114$_{388\text{-}428}$ and Mei4$_N$ at pH 7.4 (magenta), 6.9 (blue) and 6.5 (gold). Additional peaks, highlighted in dashed boxes, appear at the lower pH values. Samples with varied pH values were prepared in 25 mM $NaHPO_4$, 100 mM NaCl, 0.5 mM EDTA, 1 mM TCEP, 0.05% $NaN_3$, 5% $D_2O$. Spectra were collected at 25 °C at 500 MHz ($^1$H, Weill Cornell Medicine).

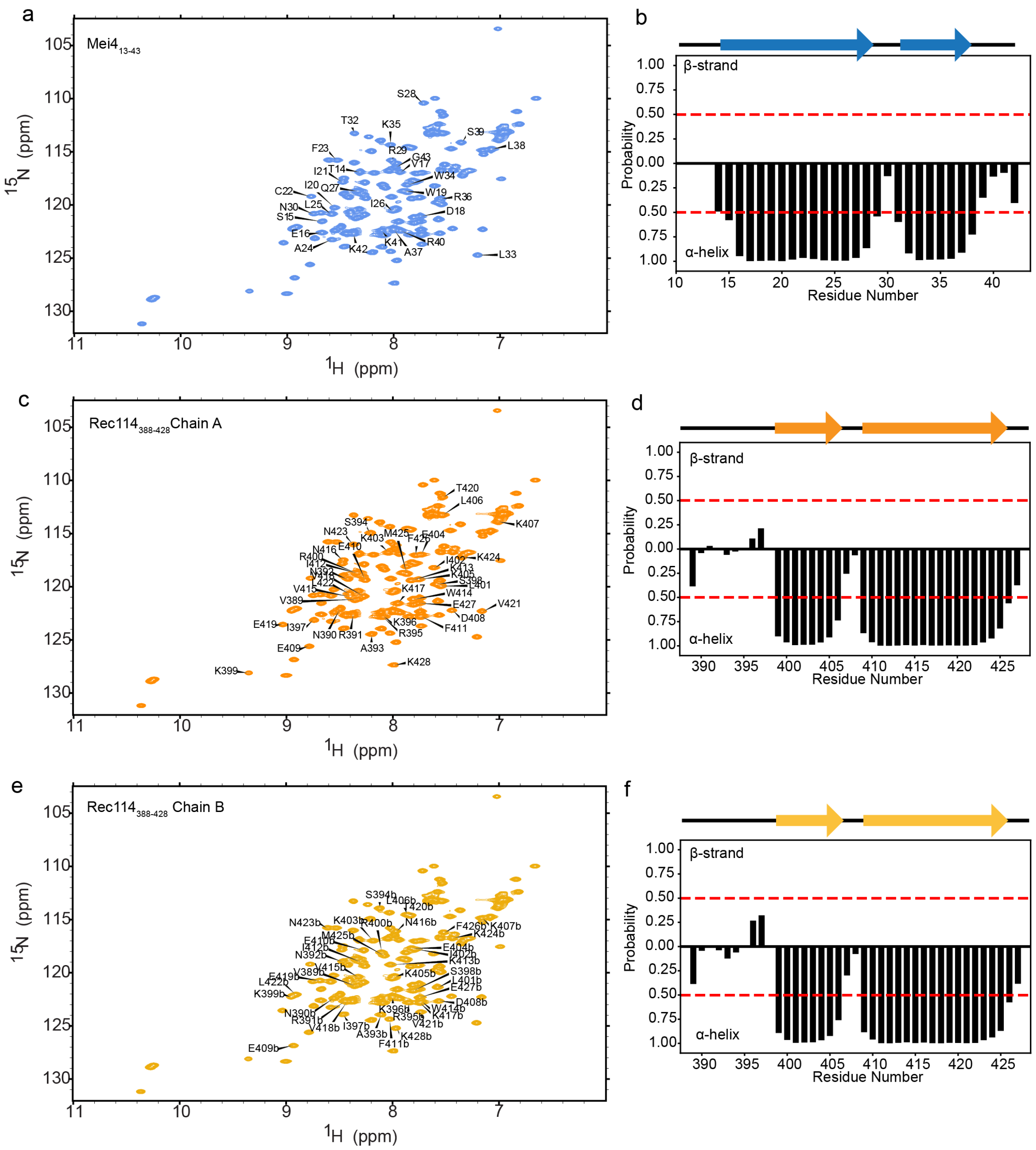


**Fig. s4: NMR signals revealed at pH 6.1 indicate that the N-terminal region of $Rec114_C$ is not structured. (a, c, e)** Assigned amide resonances for $Mei4_{13\text{-}43}$ (panel a) and the two $Rec114_{388\text{-}428}$ chains (panels c and e) within the $Rec114_{388\text{-}428}$–$Mei4_{13\text{-}43}$ complex. (**b, d, f**) TALOS-N secondary structure assessment for these residues. Stable helical structure was retained with the N-terminal truncation of Mei4. For both Rec114 chains, the previously seen helical structure was retained in this shorter construct, but the newly assigned residues 389–399 lack regular secondary structure.

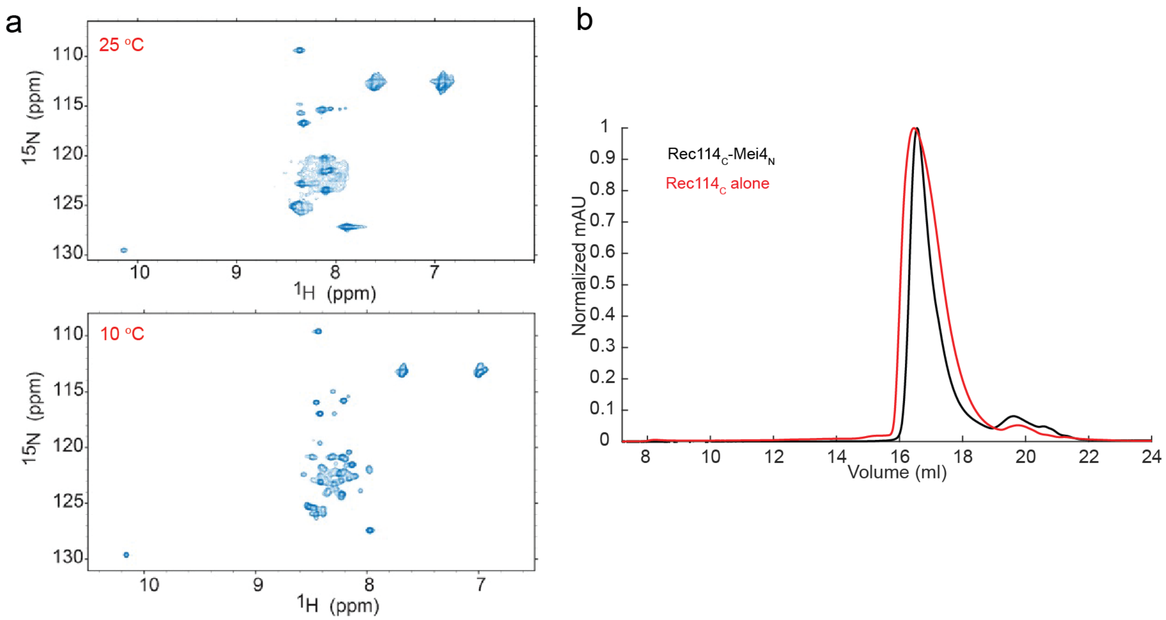


**Fig. s5: Dimers of Rec114$_C$ alone are poorly structured. (a)** Two-dimensional {$^1$H-$^{15}$N} HSQC spectra of Rec114$_C$ alone at 25 °C (top) and at 10 °C (bottom) show substantially less signal dispersion than for the trimeric complex, suggesting that Rec114$_C$ alone is substantially unfolded and/or aggregated. **(b)** Size exclusion chromatography comparing Rec114$_C$ alone with Rec114$_C$ plus Mei4$_N$.

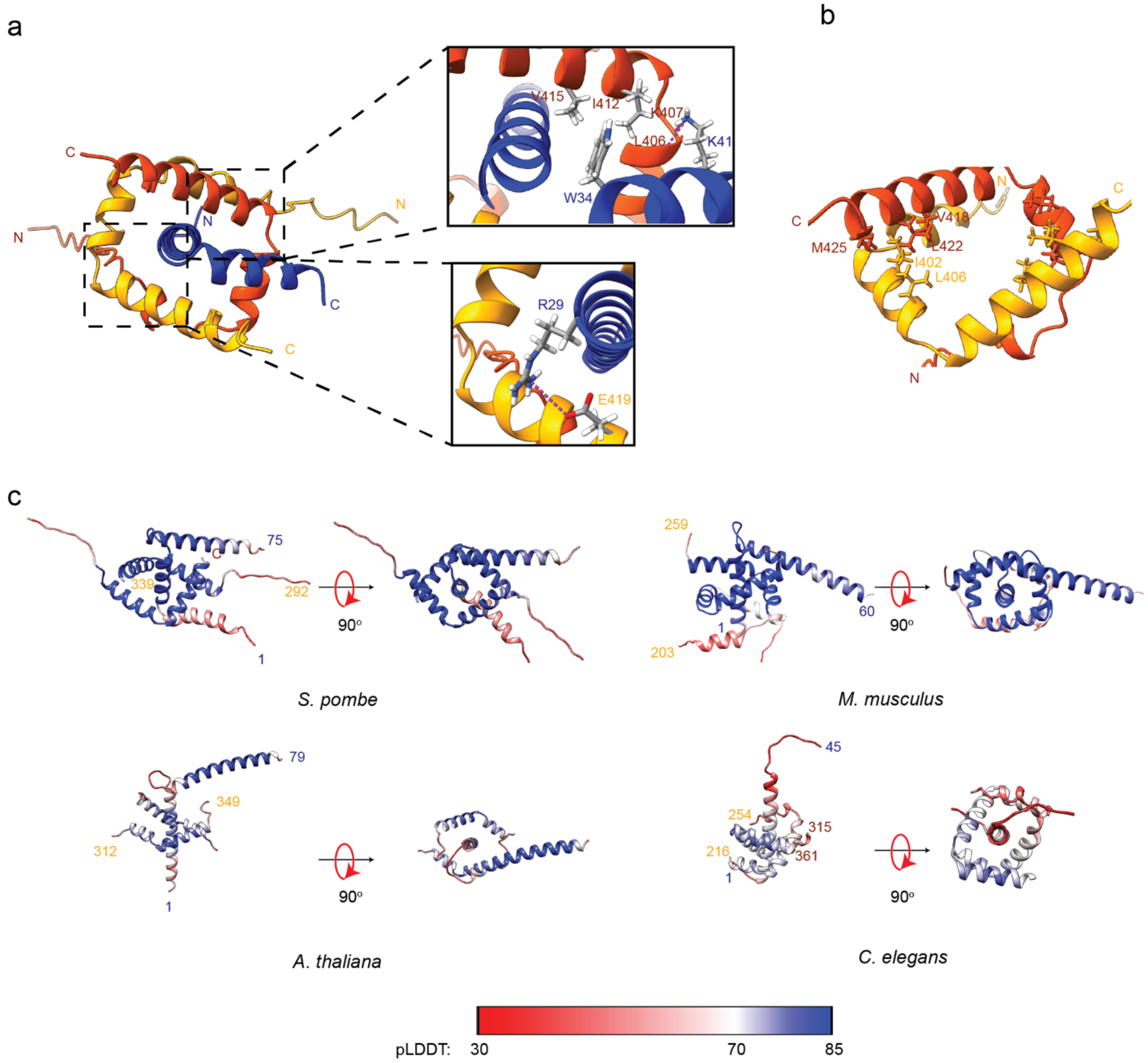


**Fig. s6: Structure and conservation of Rec114–Mei4 interactions. (a)** Asymmetric interactions between Mei4$_N$ and one chain of Rec114$_C$. The upper zoomed detail highlights hydrophobic contacts between W34 in Mei4 $\alpha$-helix 2 and I412 and V415 in $\alpha$-helix 3 from one copy of Rec114. Also shown are predicted hydrogen bonds between K41 in Mei4 $\alpha$-helix 2 and the carbonyl oxygens of L406 and K407 from one copy of Rec114. The lower zoomed detail highlights a predicted salt bridge between Mei4 R29 and E419 from one copy of Rec114. **(b)** Hydrophobic interfaces between the two copies of Rec114. **(c)** AlphaFold2 models for orthologous Rec114$_C$–Mei4$_N$ complexes from the indicated species from **Fig. 3d**, color coded by pLDDT score. The starting and ending residue numbers are indicated.

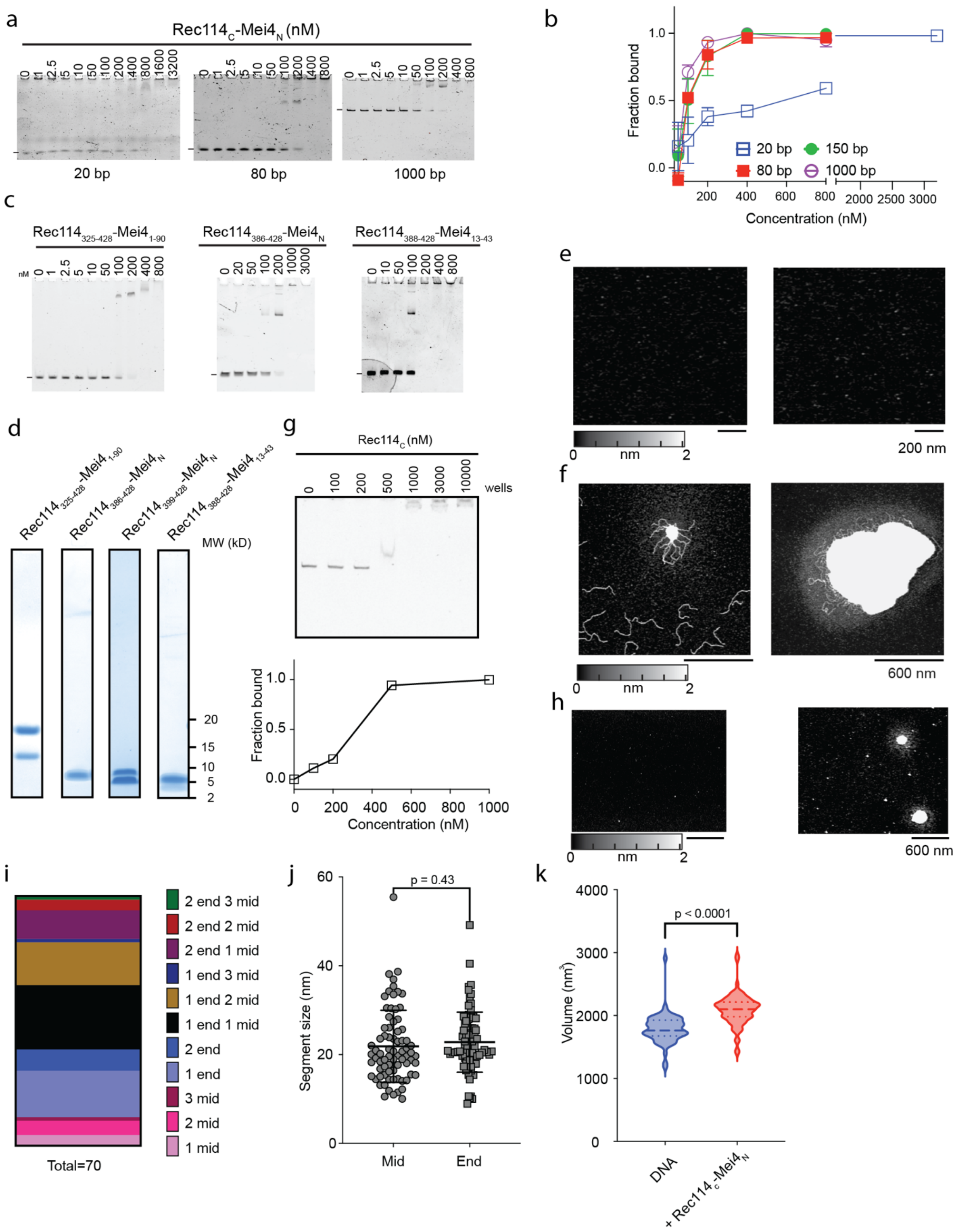


**Fig. s7: DNA binding properties of the RM-TDB domain. (a, b)** EMSAs testing binding of the RM-TDB domain to DNA substrates of varying lengths. Error bars indicate mean ± range (from two replicate experiments) or mean ± SD (from three replicates). Apparent affinities for the different DNA lengths were: 20 bp, 600 ± 100 nM (N = 2 experiments); 80 bp, 100 ± 20 nM (N = 2); 150 bp, 90 ± 30 nM (N = 3, reproduced from **Fig. 4b**); 1000 bp, 80 ± 20 nM (N = 2). **(c)** EMSAs testing binding to a 150-bp DNA substrate for different RM-TDB constructs. Quantification is in **Fig. 4b**. **(d)** SDS PAGE of purified protein complexes of varying lengths. **(e)** Representative AFM images of 2 µM RM-TDB domain in the absence of DNA. **(f)** Representative AFM images of condensates formed with 1 ng/µl 1000-bp DNA by $Rec114_{388\text{-}428}$ complexes with $Mei4_N$. Left, 200 nM protein concentration; right, 1 µM. **(g)** EMSA of $Rec114_C$ alone binding to 1000-bp DNA and its quantification (right). This experiment was conducted once; the apparent $K_d$ is ~300 nM. **(h)** Representative AFM images of 6 µM $Rec114_C$ in the absence (left) or presence (right) of 1.7 ng/µl supercoiled pUC19 plasmid DNA. **(i)** Complexes of $Rec114_C$–$Mei4_N$ (70 nM) bound to 1 ng/µl 1000-bp DNA substrate, categorized by the number and position of the DNA-bound protein particles at either the DNA end or internally (mid). **(j)** Lengths of individual protein-bound DNA segments, grouped according to position at the DNA end (N = 76) or internally (mid, N = 80). Lines represent mean ± SD. **(k)** Comparison of total DNA or protein-DNA volumes in the absence (blue, N = 63) and presence (red, N = 80) of $Rec114_C$–$Mei4_N$. Median and quartiles are indicated by dashed and dotted lines, respectively. The p values in panels j and k are from unpaired two-tailed Student's t-tests.

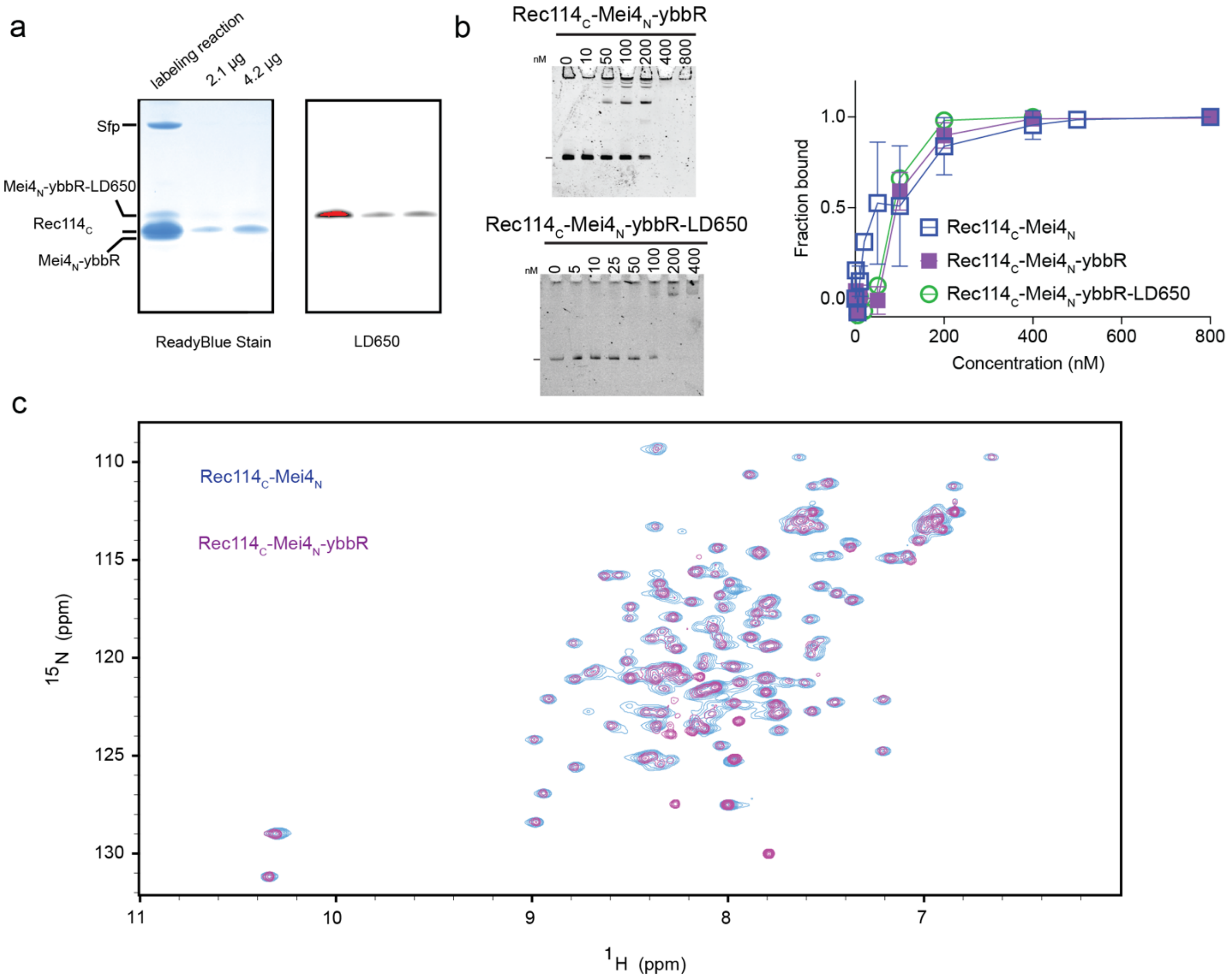


**Fig. s8: Fluorescent labeling of the RM-TDB domain. (a)** Purified complexes of Rec114$_C$ plus Mei4$_N$-ybbR were covalently labeled with LD650 using Sfp and analyzed by SDS-PAGE followed by ReadyBlue staining (left) or fluorescence scanning (right). After labeling, the sample was further purified by SEC to remove Sfp and unincorporated dye. First lane: labeling reaction; second and third lanes: SEC-purified protein. Mei4-ybbR covalently labeled with LD650 showed a shift to lower mobility. **(b)** EMSA comparing ability of the RM-TDB domain with or without the ybbR tag to bind to a 150-bp DNA substrate; the tagged protein was also tested before and after the fluorescent labeling reaction. Error bars indicate mean ± range for two different experiments with the ybbR-tagged protein (apparent $K_d$ of 90 ± 20 nM), or mean ± SD from three replicates for untagged (reproduced from **Fig. 4b** to aid comparison). The EMSA was conducted once for the labeled protein preparation (apparent $K_d$ of ~90 nM). **(c)** Comparison of the {$^1$H-$^{15}$N} HSQC spectrum of RM-TDB carrying the unlabeled ybbR tag with that of untagged RM-TDB shows additional signals, likely originating from the ybbR peptide, but otherwise no significant changes.

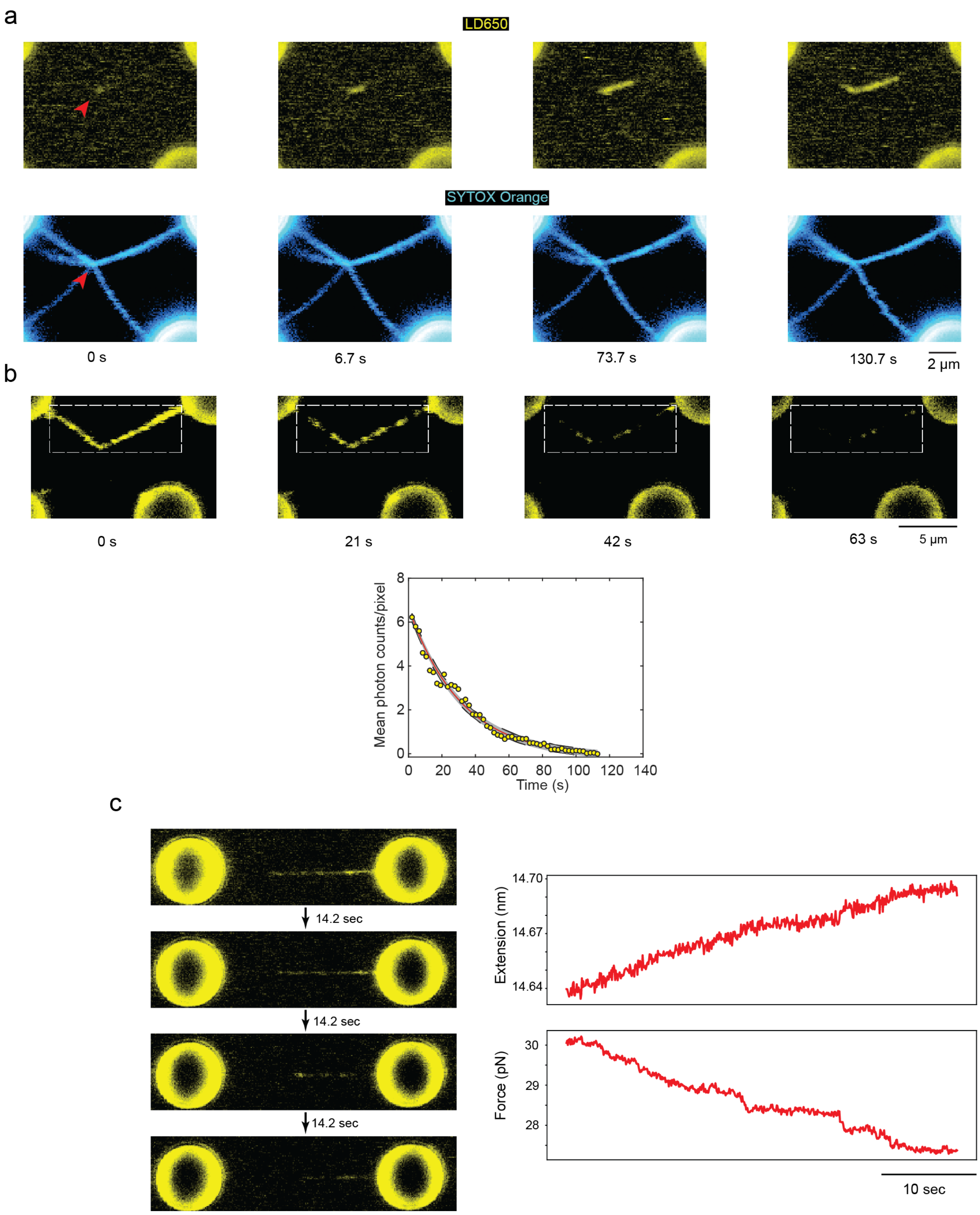


**Fig. s9: DNA bridging by the RM-TDB domain. (a)** Preferential initiation of RM-TDB binding at a crossing point. A quadruple-trap setup was used to capture two tethered pairs of beads. The top pair is connected by multiple λ DNA tethers, while the bottom pair is connected by only a single DNA. The two sets of tethers were wrapped around each other to form a single crossing point, then moved into the protein channel to allow RM-TDB to bind. The protein bound initially at or near the crossing point (red arrow), then spread over time to the regions where DNA could be coaligned. **(b)** Reversibility of RM-TDB bridges. To quantify the dissociation rate when beads with preassembled bridge-tether assemblies were moved to the protein-free channel, a region of interest (ROI) was drawn as indicated by the white dashed box for all the frames in the movie. The mean photon count per pixel was then determined for each frame and plotted as a function of time (below). The data was then fit to a single-exponential curve. For this example, the dissociation rate was 0.032 $s^{-1}$ (95% confidence interval: [0.031 $s^{-1}$, 0.033 $s^{-1}$]). See also Methods. **(c)** Force-dependent reversal of RM-TDB bridges. In contrast to the experiment shown in **Fig. 5f,** here the optical traps were held in the protein-containing channel at a fixed distance apart that placed an initial force of ~30 pN on a preassembled bridge-tether assembly. Dissociation of the RM-TDB domain was accompanied by an increase in the DNA extension (distance between the beads themselves) and a concomitant decrease of the force.

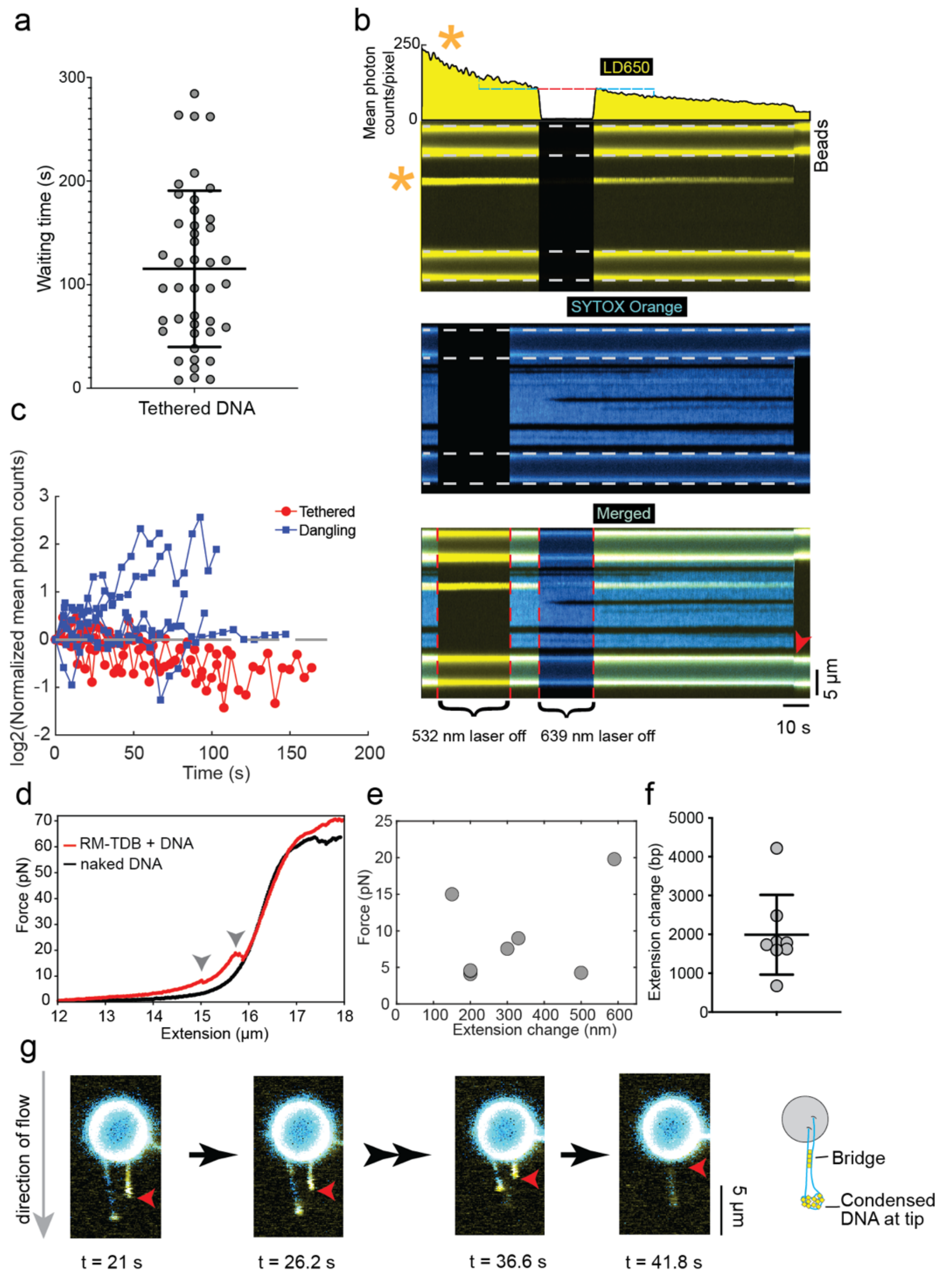


**Fig. s10: Characterization of focal DNA binding by the RM-TDB domain. (a)** Waiting times for appearance of foci of the RM-TDB domain binding to stretched DNA tethers. Each point shows the time from transfer of the bead-bound DNA into the protein channel to the time when a focal binding event was first detected. Error bars are mean ± SD. **(b)** Kymograph showing a representative RM-TDB focus (asterisk) stably bound to a stretched DNA tether, held in the fluorescent protein-containing channel. The bead boundaries are indicated by white dashed lines. Each laser was turned off briefly to show that there is negligible crosstalk between the DNA and protein fluorescence channels. Double-stranded DNA regions were stained by SYTOX Orange (shown in cyan); the dark regions are single-stranded regions that formed when duplex DNA next to spontaneous nicks was denatured by the 60 pN force applied. The RM-TDB focus remained stably bound to the tether with little or no change in position until the tether broke (indicated by the red arrow at the end). The graph above the image for the LD650 channel shows the decrease in fluorescence intensity over time for the focal binding event shown in the kymograph. Because each labeled RM-TDB trimer has a single fluorophore, the smooth decay in fluorescence signal without obvious individual steps indicates that many copies of the RM-TDB domain are present in a single focus. The intensity immediately before and after the time when the 639 nm laser was off (red dashed line) was essentially identical. This contrasts with the decrease seen during illumination periods of the same duration (blue dashed lines), therefore the signal decay (~0.01 $s^{-1}$ (range 0.007–0.013 $s^{-1}$, from fitting before and after 639 nm laser off signal decay)) is primarily due to photo bleaching. **(c)** Change in protein fluorescence intensity over time for focal binding events on stretched tethers (red, N = 4) or on dangling DNA (blue, N = 9). Each trace represents one binding event. These are the individual traces that are averaged in **Fig. 5h**. The protein fluorescence intensity at each time point was normalized to the signal in the first frame where binding of RM-TDB was detected (see Methods). **(d,e,f)** Evidence for force-dependent disruption of long-range interactions between distal DNA segments bound by RM-TDB foci. Panel d compares force-extension curves between naked and protein-bound λ DNA. Arrows indicate examples of abrupt transitions in the curve for protein-bound DNA. Panel e shows the variation of applied force and DNA extension distance for abrupt transitions in force-extension curves from 6 different molecules. Panel f shows the distribution of estimated DNA extension lengths from the transitions in panel e. Lines indicate mean ± SD. The disruption forces ranged from 4–20 pN, with an average disrupted DNA length of ~2000 bp. (**g**) A representative example showing stretches of RM-TDB binding on the coaligned arms of a dangling λ DNA molecule coincident with brighter signal at the tip (red arrow). Over time, the DNA was pulled upward against flow toward the bead surface. A cartoon is provided to aid illustration of the process on the right.

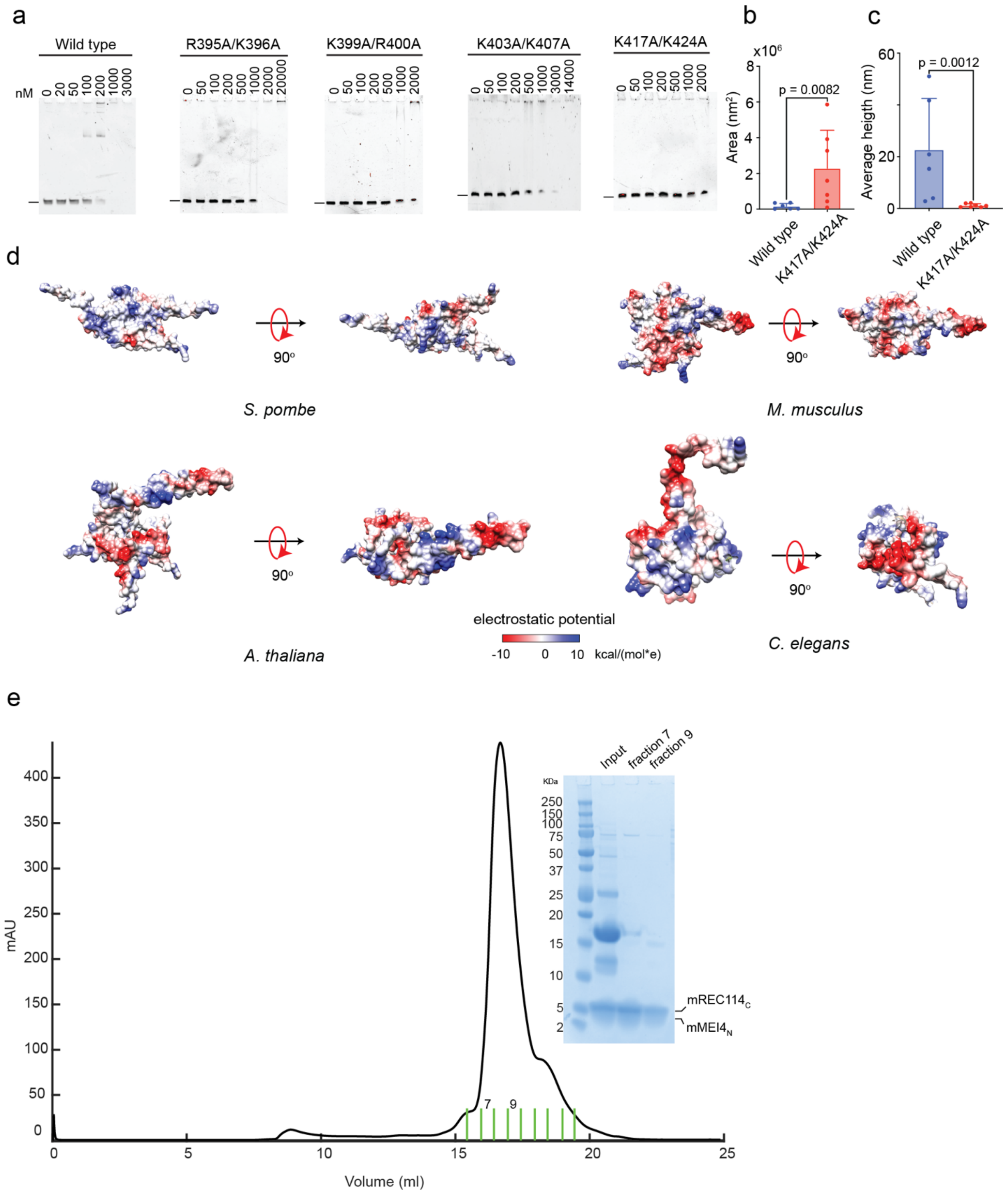


**Fig. s11: Conservation and variation of DNA-binding properties of the RM-TDB domain.** **(a)** Representative EMSAs of wild-type and mutant RM-TDB domains (complexes of $Rec114_{388-428}$ with $Mei4_N$) binding to a 150-bp DNA substrate. Quantification is in **Fig. 6b**. **(b, c)** Comparison of the distributions of areas (panel b) and heights (panel c) of condensates formed by either wild type (N = 6 condensates) or the K417A/K424A mutant (N = 7) measured by AFM as in **Fig. 6c**. Error bars indicate SD. The p values are from two-sided Mann-Whitney tests. **(d)**

Electrostatic surface potentials for orthologous Rec114$_C$–Mei4$_N$ complexes from the indicated species. Views are the same as in **Fig. s6c**. (**e**) SEC purification of mouse RM-TDB domain (mREC114$_C$-MEI4$_N$). The input to the column and fractions 7 and 9 were analyzed by SDS-PAGE and ReadyBlue staining.
